# Requirement for oxidation of neuronal ketone bodies in aging and neurodegeneration

**DOI:** 10.64898/2026.03.24.712448

**Authors:** Joyce Yang, Mitsunori Nomura, Jonathan X Meng, Thelma Y Garcia, Timothy R Matsuura, Daniel P Kelly, Ken Nakamura, John C Newman

## Abstract

Glucose is the brain’s primary fuel, but the brain can also use alternative energy substrates, especially during development or starvation. Emerging evidence suggests ketone metabolism may help the brain adapt to energy stress in neurodegenerative diseases such as Alzheimer’s disease, although its role in constitutive brain function in normal aging is poorly understood. Using iPSC-derived human neurons and adult-inducible, neuron-specific *Bdh1* knockout mice, we show that ketone body metabolism is essential for maximum energy production, neuronal function, and mouse survival—even under normal nutritional conditions. Mechanistically, phenotypes of *Bdh1* knockout neurons are mitigated by provision of acetoacetate, a downstream energy metabolite. Moreover, loss of neuronal ketone oxidation markedly increases mortality and memory deficits in Alzheimer’s disease model mice. These findings identify ketones as critical neuronal fuels, with particular importance during neurodegeneration. While non-energetic activities of ketone bodies are increasingly appreciated, oxidation for energy provision is an essential mechanism for normal function in neurons and mice. Targeting the energetic function of ketones may thus offer new therapeutic strategies for both aging and neurodegenerative diseases such as Alzheimer’s.

## Main Text

Glucose is the primary oxidative fuel for the mammalian brain under physiologic conditions (*1*). However, in energetically compromised states, the brain can rewire to utilize alternative substrates including ketone bodies, which can support the majority of energetic needs during starvation (*2*). Little is known, however, about neuronal requirements for ketone metabolism under non-fasting physiological or pathological conditions in the adult brain (*3*).

The ketone bodies β-hydroxybutyrate (βHB) and acetoacetate (AcAc) are small molecules synthesized primarily in the liver from lipid substrates. These ketones circulate and provide a readily oxidizable source of Acetyl-CoA and ATP to meet the energetic needs of diverse tissues (*4*). βHB and AcAc are interconverted by the bidirectional enzyme β-hydroxybutyrate dehydrogenase 1 (Bdh1) as the final step in ketone body production and the first step in oxidation (fig. 1A). While blood levels of ketone bodies are induced 10-to 100-fold under glucose-deprived states such as fasting, starvation, or the low-carbohydrate ketogenic diet, a small amount of ketone bodies are produced constitutively, resulting in blood levels in the 50-100 micromolar range (*5*). Astrocytes may also generate βHB and AcAc from fatty acids *in situ* (*6, 7*), a process required in *Drosophila* for normal memory function (*8*). While the role of ketones in supporting the mammalian brain during starvation is well known, their function under normal conditions is less understood.

**Fig. 1.**
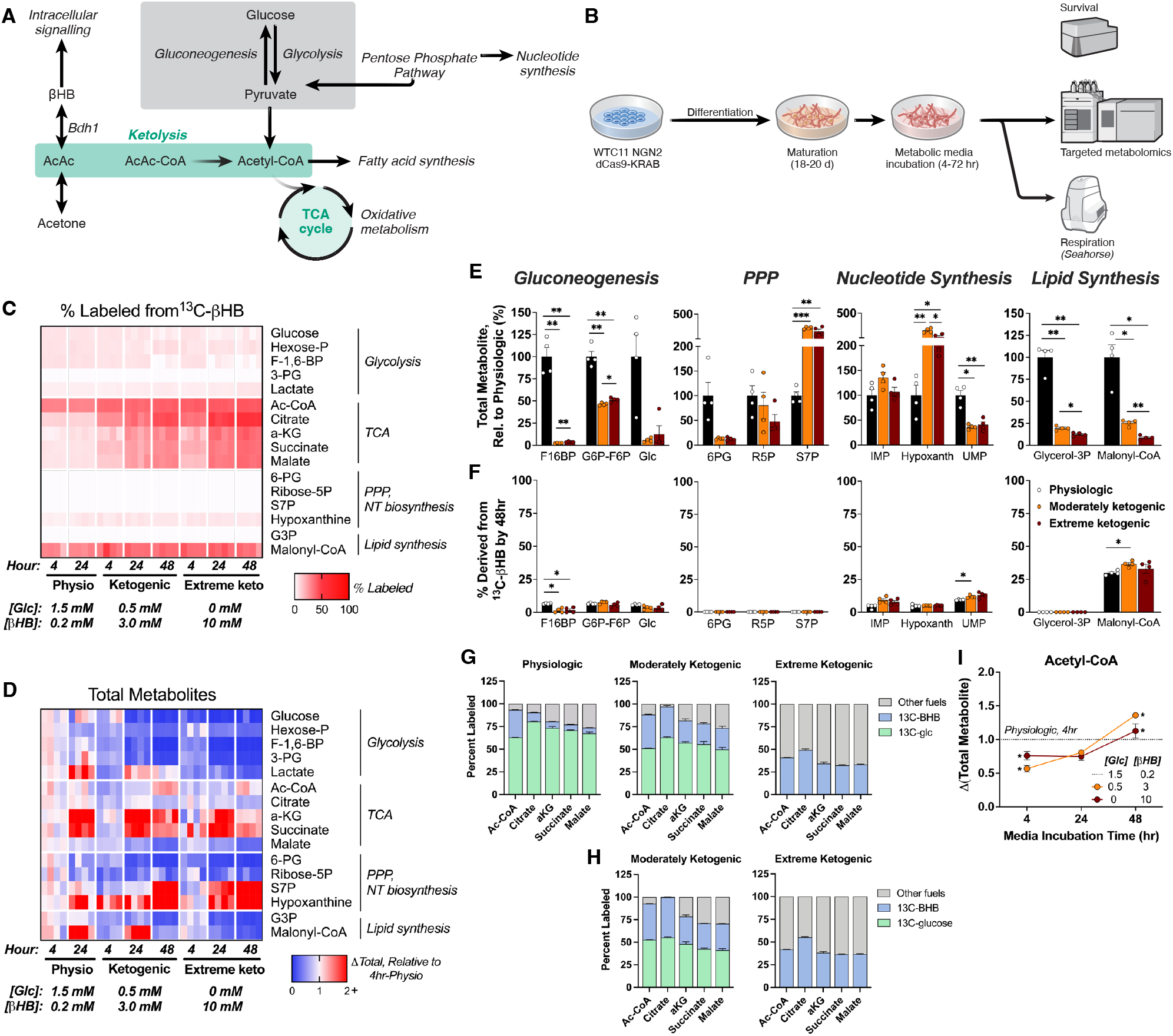
Neurons use β-hydroxybutyrate in oxidative metabolism under physiologic and ketogenic conditions. **(A)** Pathways of interest relative to ketones β-hydroxybutyrate (βHB), acetoacetate (AcAc), and acetone. **(B)** Experimental timeline for WTC11 human iPSCs expressing doxycycline-inducible Neurogenin-2. **(C)** HPLC-MS targeted metabolomics conducted on 20-22d neurons incubated with ^13^C-βHB under physiologic, moderately ketogenic, or extreme conditions for 4, 24, or 48h. *Rows*: metabolites; *columns*: percent labeled from ^13^C-βHB; and **(D)** total metabolite change per replicate, normalized to 4hr physiologic average. N=4 per condition. **(E)** Physiologic-normalized total metabolite and **(F)** percent derivation from ^13^C-βHB after 48hr incubation in metabolic media. N=4 per condition. **(G)** Fractional labeling of TCA metabolites from isotope tracing of ^13^C-glc or ^13^C-βHB after 24hr or **(H)** 48hr incubation in ketogenic conditions. N=4, except 24 h “moderately ketogenic” with ^13^C-glc (3). **(I)** Changes in total Acetyl-CoA over time, relative to 4hr physiologic. N=4 per condition. Data are means ± SEM. *p < 0.05, **p < 0.01, and ***p < 0.001 by Brown-Forsythe and Welch one-way ANOVA with Dunnett’s T3 multiple comparisons (E-F), or 2-way ANOVA with Dunnett’s multiple comparisons vs. physiologic 4hr average (I).

Ketogenic metabolism is required in the neonatal state for normal mammalian brain and sensory neuron development (*9, 10*), but less is known about its requirement beyond this period. Glucose and energy utilization in the brain decreases with age and in Alzheimer’s disease (AD) (*11-15*), as well as in cognitively normal ApoE4 carriers (*16, 17*) who are at increased risk for AD. Baseline brain usage ratios of 29:1 for glucose to other fuels can shift as low as 2:1 in incipient AD (*18, 19*). The capacity to oxidize ketone bodies remains intact in AD (*20*), and in heart failure, cardiomyocytes preferentially utilize ketone bodies as an adaptive means to rescue energy deficits (*21, 22*). Ketogenic therapies are therefore under clinical investigation in both heart failure and AD (*12, 21*). Data on ketone body utilization in AD is largely derived from exogenous administration or ketogenic diets (*23, 24*), but not the study of endogenous constitutive ketogenesis. It is unknown whether neuronal metabolism of endogenous ketone bodies is compensatory for bioenergetic deficits in AD, or more broadly, whether this process is required for normal neuronal and brain function throughout adult life.

Here we use a combination of induced pluripotent stem cell (iPSC)-derived human neurons and adult-inducible, neuron-specific mouse knockouts of Bdh1 to interrogate the role of ketone bodies in neuronal metabolism and organismal aging, in both wild-type and AD genetic backgrounds. Using our models, we find a neuronal requirement for ketone bodies to generate maximal acetyl-CoA, with distinct roles for AcAc and βHB. Further, mice with adult-induced neuron-specific Bdh1 knockout show accelerated early mortality and memory impairments in both wild-type and AD genetic backgrounds, supporting a constitutive requirement for neuronal ketone body oxidation throughout mammalian adulthood.

## Results

### Human neurons rapidly rewire metabolic pathways in response to ketogenic environments

Although β-hydroxybutyrate (βHB) is constitutively present in blood, crosses the blood-brain barrier, and is oxidized in the brain (*2, 12*), there is little direct evidence of neuronal uptake and metabolism of βHB, in part because brain tissue and primary neuronal cultures contain a significant proportion of glial cells. To overcome this hurdle, we studied βHB metabolism in near-homogenous human excitatory neurons derived from iPSCs (*25, 26*) co-expressing dCas9-KRAB and neurogenin-2 (Ngn2) (*27*).

iPSC lines were differentiated into neurons by doxycycline induction of Ngn2 and cultured for 20 days (fig. 1B). Neurons were then incubated for 4, 24, or 48 h in media with different ratios of ^13^C-βHB to glucose—physiologic (1.5 mM glc + 0.2 mM βHB; based on brain microdialysis (*28-32*)), moderately ketogenic (0.5 mM glc + 3 mM βHB; simulating bloodstream levels during ketogenic diet (*33*)), or extreme ketogenic (10 mM βHB)—before metabolite isolation. Isotope tracing of ^13^C-βHB revealed rapid uptake and incorporation of βHB into the TCA cycle and lipid synthesis, but not into other major pathways (fig. 1C,F). Surprisingly, a small amount of glucose, glucose-6-phosphate (G6P), fructose-6-phosphate (F6P), and fructose-1,6-bisphosphate (F1,6BP) also contained ^13^C (fig. 1F), suggesting gluconeogenesis may be occurring in neurons.

The metabolomic profiles of cells exposed to moderate and extreme βHB/glucose ratios differed from cells exposed to a physiologic ratio (fig. 1D), as illustrated by a decreased flux through glycolytic metabolites detectable within 4 h of incubation and an increased level of some TCA metabolites by 24 h. There was also a substantial elevation in nucleotide (NT) biosynthesis, with sedoheptulose-7P and hypoxanthine appearing as two of the strongest hits across the panel (fig. 1E). Changes were similar across moderate and extreme ketogenic conditions, suggesting that large changes in fuel availability are not necessary to significantly alter the neuronal metabolome.

### Human neurons utilize βHB for oxidative metabolism under physiologic states

Labeling of TCA metabolites from ^13^C-βHB after 24 h under physiologic βHB/glucose ratio was substantial, supporting that human neurons actively utilize the constitutive ketones present in the brain microenvironment. For instance, approximately 30% of acetyl-CoA—the TCA cycle entry-point metabolite—was at least partially derived from βHB under physiologic conditions, a proportion that increased only modestly as βHB availability increased from 0.2 mM to 3 mM and 10 mM (37% and 41%, respectively) (fig. 1G, blue).

Given this substantial usage of ketones for oxidative metabolism, we wondered whether βHB is a dominant neuronal fuel source, similar to glucose. We repeated HPLC-MS targeted metabolomics with isotope tracing of ^13^C-glucose for 24 or 48 h under physiologic and moderately ketogenic conditions. Neurons in ketogenic conditions exhibited a significant shift away from glucose-based TCA derivation, and towards βHB by 24 h (fig. 1G, green vs. blue). The shift was even more pronounced by 48 h (fig. 1H). Again, large changes in fuel availability were not necessary to generate fuel shifts (physiologic vs. moderately ketogenic), suggesting that human neurons readily utilize βHB. Total Acetyl-CoA levels were also elevated in ketogenic relative to physiologic conditions by 48 h (fig. 1I). Taken together with the isotope tracing, these observations establish that human neurons directly import and metabolize βHB, and that βHB is a major fuel source contributing to neuronal oxidative metabolism across a range of conditions, including under normal glucose availability.

### AcAc is a better alternative to glucose than βHB for neuronal energy production

We next investigated whether other lipid-based fuels—ketones AcAc and acetone, and common fatty acids palmitate (PA), oleate (OA), and stearate (SA)—induce similar metabolic phenotypes as βHB does. We first generated dose-survival curves in 21-day human neurons incubated for 48 to 72 h in lipid-based media, and selected 8 mM (ketones) and 50 uM (fatty acids) as concentrations that maximized fuel availability with little or no cell death (Extended Data fig. 1A).

Using HPLC-MS targeted metabolomics, we observed decreased total and labeled glycolytic metabolites across all conditions relative to physiologic (fig. 2A), consistent with a shift to lipid-based metabolism. We performed ranked analyses on the metabolite panel for fold-change comparisons and found large elevations in inosine monophosphate (IMP) and guanosine/adenosine monophosphate (GMP, AMP) without substantial incorporation of initial ^13^C-fuels (Extended Data fig. 1B-C). Given that these metabolites are precursors of ATP, NADH, and NADPH, these peripheral elevations suggest a possible mechanism through which neurons accommodate metabolic changes by increasing coenzymes for energy production. Surprisingly, sole provision of 13C-AcAc resulted in TCA labeling similar to that of glucose under physiologic conditions (fig. 2B, “AcAc” vs. “^13^Glc+?HB”), while sole provision of ^13^C-βHB resulted in more intermediate TCA labeling similar to before (fig. 1C “extreme keto” vs. fig 2B “BHB”). This suggests that AcAc is a more effective substitute for glucose than βHB in maintaining neuronal oxidative metabolism.

**Fig. 2.**
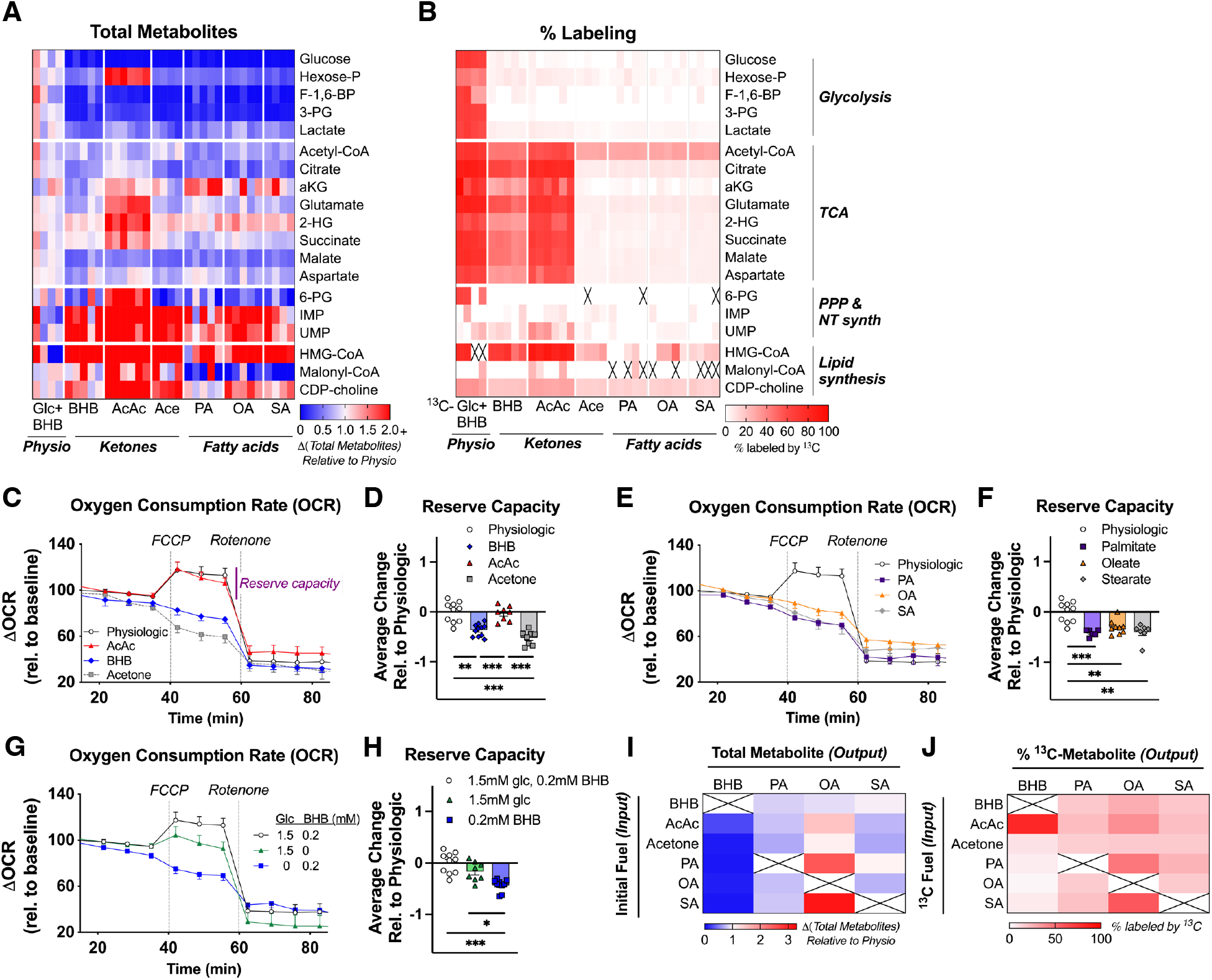
Neurons preferentially utilize acetoacetate in oxidative metabolism to support cellular respiration. **(A)** Targeted metabolomics conducted on 21d neurons after 3d incubation in physiologic, ketone-, or fatty acid (FA)-based medias with initial ^13^C-fuel of interest. *Rows*: metabolites; *columns*: total metabolite change, normalized to physiologic average; and **(B)** corresponding % labeled from ^13^C-fuel. N=4 (Glc+βHB, Ace, SA), 5 (βHB, PA, OA), or 6 (AcAc) samples/condition. **(C)** Oxygen consumption rate (OCR) after 3d incubation in ketone-based or **(E)** FA-based conditions. Cells were treated sequentially with carbonyl cyanide-p-trifluoromethoxyphenylhydrazone (FCCP) and rotenone. Data normalized to average of baseline readings per condition, highlighting differences in maximum OCR (“reserve capacity”). **(D, F)** Quantification of relative reserve capacity. N=6 (PA), 7 (SA), 8 (AcAc, Ace), 9 (OA), 10 (Physio), or 11 (BHB) samples/condition across 2 experimental replicates. **(G)** OCR changes after 3d incubation in physiologic media variants (+-glucose, +-βHB), and **(H)** quantification of reserve capacity. N=8 (Glc), 9 (βHB), 10 (Glc+βHB) samples/condition across 2 experimental replicates. **(I)** Total metabolite levels of BHB, PA, OA, and SA (*columns*) based on initial fuel (*rows*), relative to physiologic levels; and **(J)** percent of each “output” containing ^13^C from “input”. N=4 (Ace, SA), 5 (βHB, PA, OA), or 6 (AcAc) samples/condition. Data are means ± SEM. *p < 0.05, **p < 0.01, and ***p < 0.001 by Brown-Forsythe and Welch one-way ANOVA with Dunnett’s T3 multiple comparisons.

### Ketones are vital to maximize mitochondrial respiration under restricted and physiologic states

To further explore contributions of lipid-based fuels to mitochondrial oxidative phosphorylation, we recorded oxygen consumption rates (OCR) via Seahorse assay in mature neurons incubated for 72 h in the previously-described conditions. Of the sole-provided fuels, only AcAc (but not βHB) matched the reserve capacity observed under physiologic conditions containing both glucose and βHB (fig. 2C-F), supporting the hypothesis that AcAc is a more effective respiratory substrate than βHB, that can substitute for both glucose and βHB to maintain neuronal mitochondrial respiration. Given the previously demonstrated contribution of βHB to the acetyl-CoA pool under physiological conditions (fig. 1G), we next assessed the effect of physiologic fuel availability on OCR. Removal of this low concentration of βHB led to a significant decrease in reserve capacity (fig. 2G-H), underscoring the requirement for βHB during physiologic states. Furthermore, incremental changes in fuel availability produced corresponding changes in reserve capacity, suggesting that neurons are highly sensitive to small fluctuations in environmental ketone (Extended Data fig. 1D) and glucose (Extended Data fig. 1E) levels. Therefore, although βHB alone cannot compensate for loss of glucose (fig. 2C), βHB supplementation can boost energy production beyond glucose alone.

### Human neurons preferentially convert excess lipid-based fuels into oleic acid

The fuel concentrations utilized for metabolomics and Seahorse were selected to maximize availability while minimizing toxicity (Extended Data fig. 1A). To gain insight into whether neurons perform ketogenesis or store excess ketones as fatty acids, we screened interconversion patterns between each of the tested fuels from the metabolomics panel. Analyses of total metabolite production (fig. 2I) and percent derived from each starting fuel (fig. 2J) revealed the expected conversion of AcAc to βHB, but no indication of *de novo* ketogenesis, and instead preferential conversion of all substrates towards storage as oleic acid via fatty acid synthesis (fig. 1A).

### Knocking down Bdh1 causes oxidative and survival deficits in human neurons that are mitigated by AcAc supplementation

Bdh1 is the bidirectional enzyme that converts βHB into AcAc, which proceeds via Acetyl-CoA into the TCA cycle. To test the importance of this conversion for neuronal health and metabolism, we knocked down Bdh1 in iPSCs (*BDH1-kd* line) via inducible dCas9-KRAB (*27*) and differentiated the iPSCs into neurons. As controls, we generated an iPSC line expressing a non-targeting guide RNA (*NTG* line) and cultured for 21 days to validate stable knockdown (fig. 3A). Survival curves showed that incubation with either 3 mM or 8 mM βHB, but not AcAc, decreased the survival of *BDH1-kd* neurons relative to *NTG* control (Extended Data fig. 2A-B, fig. 3B). To determine if these differences may be attributed to discrepancies in energy production, we conducted targeted metabolomics in basal (unrestricted glc + pyruvate), physiologic (1.5 mM glc + 0.2 mM βHB), physiologic AcAc (1.5 mM glc + 0.2 mM AcAc), βHB-only (3 mM βHB), or AcAc-only (3 mM AcAc) medias. As expected, *BDH1-kd* neurons largely stopped metabolizing 13C-βHB (fig. 3D), resulting in reduced TCA metabolites relative to physiologic (fig. 3C). General usage of ^13^C-AcAc was unaffected (fig. 3D). *BDH1-kd* neurons incubated in βHB also exhibited reductions in S7P and dGMP compared to control, and to AcAc-incubated *BDH1-kd* and control neurons. This suggests these peripheral elevations are linked to ketone oxidation via Bdh1.

**Fig. 3.**
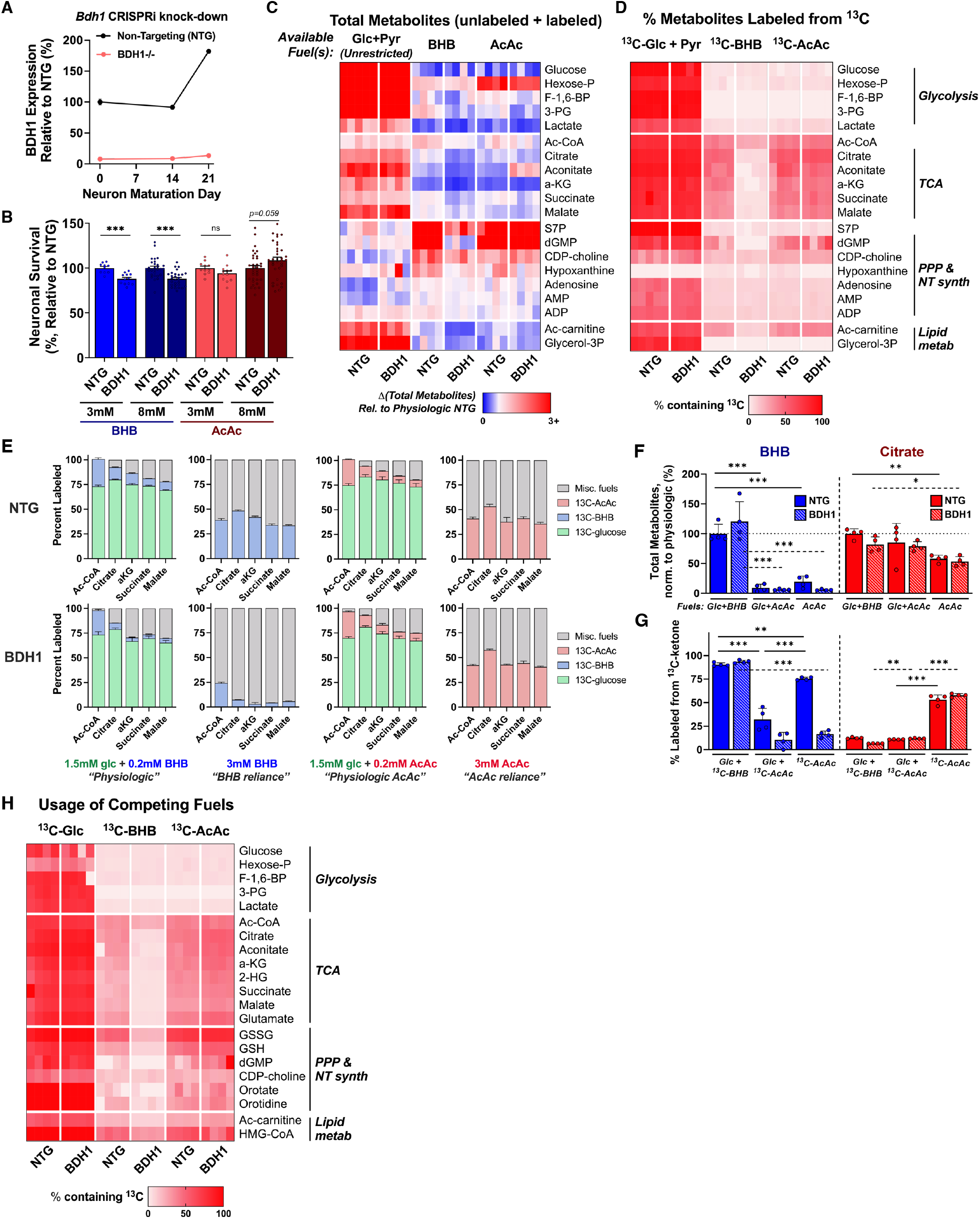
AcAc mitigates deficits as a preferential oxidative fuel in *Bdh1-KD* human neurons. **(A)** Validation of *Bdh1* knockdown (BDH1) vs. non-targeting control (NTG) via dCas9-KRAB in WTC11 neurons. **(B)** Survival of 21d BDH1 and NTG neurons after 3d incubation in βHB- or AcAc-only conditions, normalized to NTG. N=10, 12, 29, 28, 11, 12, 35, 34 (*left to right*) samples across 2 experimental replicates. Data are means ± SEM. **(C)** Targeted metabolomics after 3d incubation in basal (^13^C-Glc + Pyr, unrestricted), ^13^C-βHB-only (3mM), or ^13^C-AcAc-only (3mM) media. *Rows*: metabolites; *columns*: total metabolite change, normalized to physiologic NTG (1.5mM Glc + 0.2mM βHB); and **(D)** percent labeled from ^13^C-fuel. N=4 samples/condition, except NTG unrestricted (5). **(E)** Fractional labeling of TCA metabolites from isotope tracing of 13C-glc, -βHB, or -AcAc after 72hr. N=4 per condition. **(F)** Total levels of βHB and citrate after incubation in physiologic (1.5mM Glc + 0.2mM βHB), physiologic AcAc (1.5mM Glc + 0.2mM AcAc), or AcAc-only (3mM AcAc) conditions; and **(G)** percent of βHB or citrate containing ^13^C label from βHB or AcAc. N=4 samples/condition. Data are means ± SEM. **(H)** Metabolomics on neurons after 3d incubation in 3mM glc + 3mM βHB + 3mM AcAc. *Columns:* percent of metabolite labeled from each initial ^13^C-fuel in NTG vs. BDH1 neurons. N=4 samples/condition. *p < 0.05, **p < 0.01, and ***p < 0.001 by unpaired two-tailed t-test with Welch’s correction (B), or 2-way ANOVA with Tukey’s multiple comparisons; select comparisons shown (F-G).

To explore the possibility that AcAc mitigates Bdh1-related oxidative deficits, we compared the generation of TCA metabolites in physiologic conditions relative to either 0.2 mM βHB or 0.2 mM AcAc, and from either ^13^C-glc or ^13^C-βHB/-AcAc. In physiologic states, the relative contribution of ^13^C-βHB was reduced in *BDH1-kd* neurons (fig. 3E). These differences were mitigated in physiologic AcAc conditions, where βHB was swapped for AcAc. We observed similar mitigation when comparing ^13^C-βHB versus ^13^C-AcAc reliance conditions.

### Bdh1 strongly favors ketolysis in human neurons

In extrahepatic tissues, the canonical function of Bdh1 is ketolysis, the conversion of imported βHB to AcAc as the first step towards oxidation of βHB. But this reaction is reversible, and its direction is determined by substrate concentration gradients and NAD^+^/NADH balance. Since βHB is now known to be a multifaceted signaling metabolite (*4*), the reverse reaction could enhance these signaling activities (fig. 1A). To test the directionality of Bdh1 in neurons, we analyzed βHB and citrate production as readouts for the reverse (βHB-producing) and canonical (AcAc-generating) directions, respectively. In *NTG* neurons, only citrate was substantially produced from ^13^C-AcAc (fig. 3F-G), supporting the primacy of the canonical reaction in neurons. Moreover, the fraction of every metabolite in the panel derived from ^13^C-AcAc was nearly identical in *NTG* and *BDH1-kd* (Extended Data fig. 2D; y = 1.04x -0.14, R^2^ = 0.99), indicating no impact of Bdh1 activity on ^13^C-AcAc metabolic fate.

To probe the impact of Bdh1 on the relative usage of energetic metabolites, we conducted targeted metabolomics on 21-day *NTG* and *BDH1-kd* neurons incubated with all three substrates (βHB, AcAc, and glucose) at 3 mM each, run in triplicate with alternating ^13^C-labeled fuels. ^13^C-glc labeled a high fraction of most metabolites in our panel in *NTG* neurons, as expected, whereas 13C-AcAc and ^13^C-βHB labeled a subset of the panel and less strongly than ^13^C-glc (fig. 3H). *BDH1-kd* substantially reduced labeling by ^13^C-βHB only, consistent with the metabolic fate of βHB being primarily via conversion to AcAc through Bdh1. There was a corresponding small relative increase in labeling from *both* ^13^C-glc and -AcAc in response to *BDH1-kd*. Altogether, these data are consistent with Bdh1 functioning primarily in the canonical direction in neurons, and with neurons actively metabolizing ketones even in the presence of abundant glucose.

### Knockout of Bdh1 in neurons exacerbates mortality and memory impairments in normal aging mice

Given the high rate of neuronal ketone oxidation *in vitro*, even in the presence of abundant glucose, we hypothesized that neuronal oxidation of βHB may be required for maintenance of normal cognitive function *in vivo*, even under normal diet and housing conditions (*ad libitum* feeding, typical diet) (*34*). To test this hypothesis, we generated neuron-specific *Bdh1* knockout (KO) (Bdh1^Thy1−/−^) mice using a Bdh1^fl/fl^ line (*22*) and the Thy1-Cre^ERT2^ (SLICK-H) line (*35*). First, we generated a preliminary cohort for validation and to test acute effects of the knockout, injecting two-month-old Bdh1^+/+^ (Bdh1^fl/fl^ without Thy1-Cre^ERT2^) and Bdh1^Thy1−/−^ mice with tamoxifen. We validated knockout of *Bdh1* in NeuN-positive neurons, but not GFAP-positive astrocytes (Extended Data fig. 3, A and B). The deletion of *Bdh1* in neurons did not result in any significant alterations in *ad libitum* or fasted body weight, blood glucose levels, or plasma βHB levels (Extended Data fig. 3C). *Bdh1* KO did not alter expression of ketogenic or ketolytic genes in the brain, heart or liver, except for *Bdh1* expression in the brain (Extended Data fig. 3D). Glucose tolerance and insulin tolerance, assessed at nine months of age, were also unaltered (Extended Data fig. 4, A and B). The KO did not alter the effect of a ketogenic diet (KD) on body weight, blood glucose levels, plasma βHB levels, or the expression of ketogenic and ketolytic genes in the brain other than *Bdh1* (Extended Data fig. 4, C and D). These findings indicate that KO of *Bdh1* in neurons does not generate acute toxicity, alter systemic glucose or ketone metabolism, or alter ketone-related transcription in the short term.

We then tested the role of neuronal ketolysis in lifespan and healthspan in well-powered mouse cohorts which were administered Tamoxifen at four months of age. Surprisingly, Bdh1^Thy1−/−^ mice of both sexes had decreased survival and reduced median lifespan, with a stronger negative effect in females (Fig. 4A). At 24 months of age, Bdh1^Thy1−/−^ mice also displayed reduced memory performance compared to controls in the novel object recognition (NOR) test (Fig. 4B), as well as impaired spatial working memory in the Y-maze (Extended Data fig. 5C). There were no differences in body weight (Extended Data fig. 5A) or in other aspects of behavior or physical performance, as assessed by the open field (Extended Data fig. 5B), elevated plus maze (EPM) (Extended Data fig. 5D), and rotarod (Extended Data fig. 5E).

**Fig. 4.**
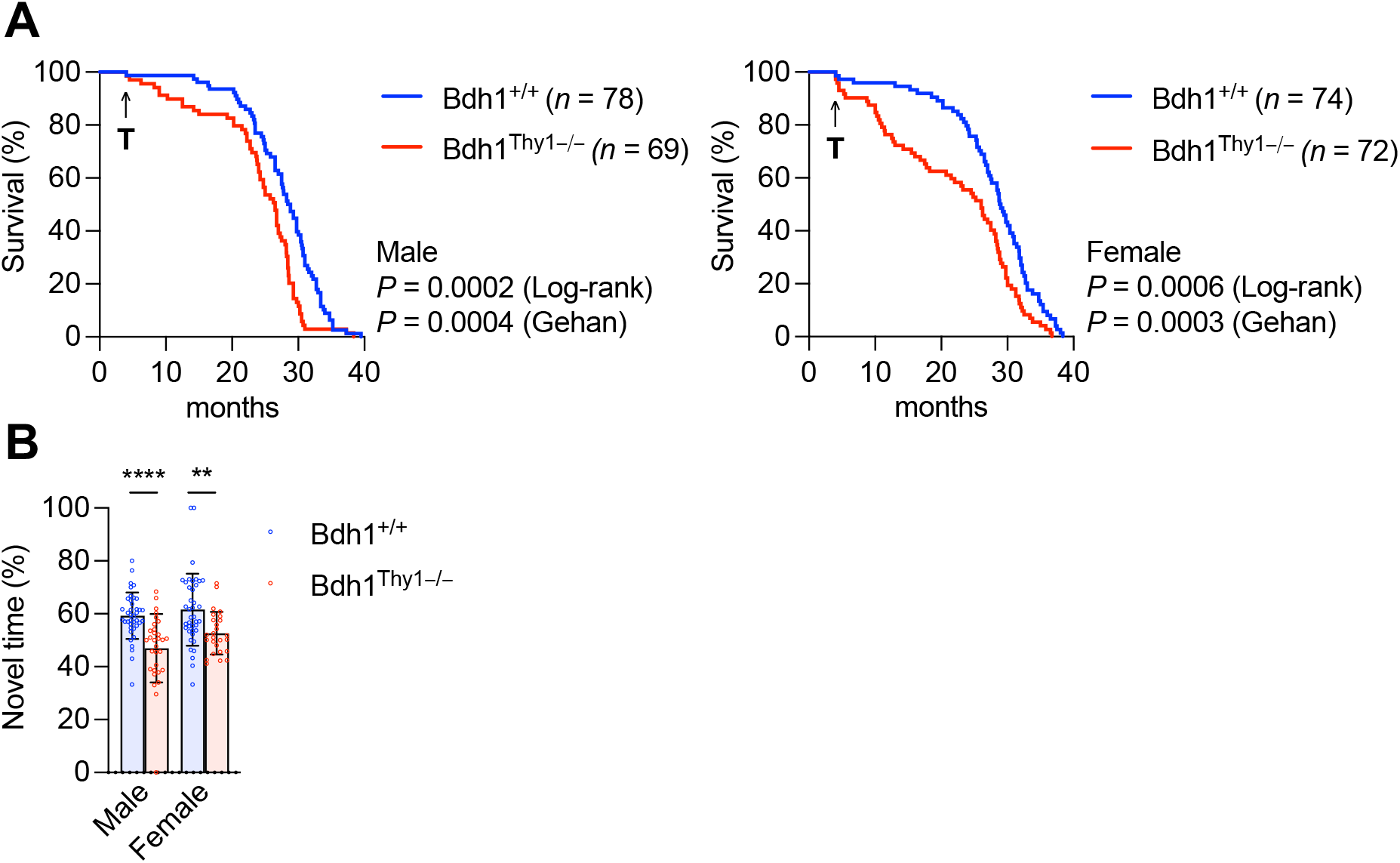
Knockout of *Bdh1* in neurons exacerbates mortality and memory impairments in normal aging mice. (A) Survival curves. Log-rank (Mantel-Cox) and Gehan-Breslow-Wilcoxon tests were used to calculate p-values. (B) Novel object recognition test at 24 months old. N=39 (Bdh1^+/+^ male), 31 (Bdh1^Thy1−/−^ male), 39 (Bdh1^+/+^ female), and 26 (Bdh1^Thy1−/−^ female) mice per group. P <0.0001 (male, t = 4.72, df = 68) and p = 0.0036 (female, t = 3.02, df = 64) by unpaired two-tailed t-test assuming Gaussian distribution. Data are means ± SD. Mice were given tamoxifen at four months old to knockout *Bdh1* in neurons.

### Knockout of Bdh1 in neurons exacerbates mortality and memory defects in hAPPJ20 mice

Finally, we hypothesized that the deleterious effects of neuronal Bdh1 deletion would be exacerbated and become apparent at younger ages in AD models which are characterized by energetic stress (*36-38*). We crossed Bdh1^Thy1−/−^ mice with hAPPJ20 AD model mice, which overexpress a human APP gene containing several disease-associated mutations (*39*), and administered tamoxifen at 4 months of age. The knockout (hAPPJ20/Bdh1^Thy1−/−^) demonstrated dramatically reduced survival, with males more strongly affected (Fig. 5A). Body weights were similar (Extended Data fig. 6A). At 12-14 months of age we observed more significant memory impairment (NOR) (Fig. 5B), margin distance (open field) (Extended Data fig. 6B), and anti-anxiety behavior (EPM) (Extended Data fig. 6D) in the female KO mice compared to hAPPJ20/Bdh1^+/+^. At 19-22 months of age, long-term memory in the Barnes maze was impaired in female hAPPJ20/Bdh1^Thy1−/−^ mice, but not in males (Fig. 5C). Conversely, in a simultaneously tested cohort on wild-type background, impaired memory in Barnes maze was observed only in male KO mice (Fig. 5C). There were no differences in tests of spatial working memory (Y-maze) (Extended Data fig. 6C) and physical function (Rotarod) (Extended Data fig. 6E). Overall, in both WT background and, more dramatically, in the hAPPJ20 AD model, impaired neuronal metabolism of BHB was associated with substantial mortality and memory function deficits, consistent with a requirement for neuronal ketone body oxidation under normal dietary conditions in both aging and AD.

**Fig. 5.**
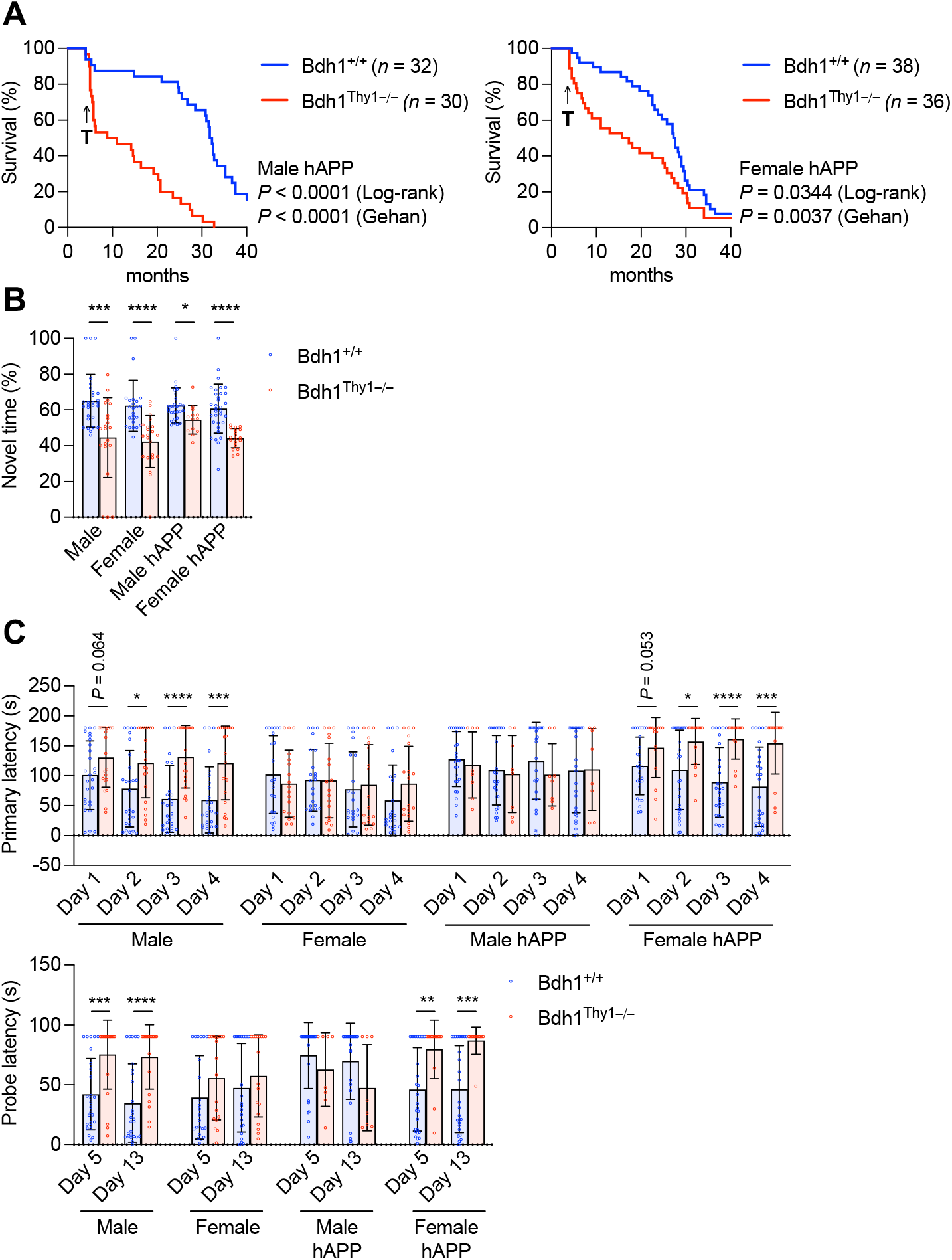
Knockout of *Bdh1* in neurons exacerbate mortality and memory impairments in hAPPJ20 mice. (A) Survival curves. Log-rank (Mantel-Cox) and Gehan-Breslow-Wilcoxon tests were used to calculate p-values. (B) Novel object recognition test at 12 months old. N=30 (Bdh1^+/+^ male), 23 (Bdh1^Thy1−/−^ male), 27 (Bdh1^+/+^ female), 22 (Bdh1^Thy1−/−^ female), 28 (hAPPJ20/Bdh1^+/+^ male), 14 (hAPPJ20/Bdh1^Thy1−/−^ male), 33 (hAPPJ20/Bdh1^+/+^ female), and 20 (hAPPJ20/Bdh1^Thy1−/−^ female) mice per group. P = 0.00033 (male, t = 3.88, df= 47), p < 0.0001 (female, t = 4.77, df = 45), p = 0.014 (hAPPJ20 male, t = 2.57, df = 39), and p < 0.0001 (hAPPJ20 female, t = 5.02, df = 50) by unpaired two-tailed t-test assuming Gaussian distribution. Data are means ± SD. (C) Barnes maze at 19-22 months old. N=27 (Bdh1^+/+^ male), 21 (Bdh1^Thy1−/−^ male), 23 (Bdh1^+/+^ female), 18 (Bdh1^Thy1−/−^ female), 27 (hAPPJ20/Bdh1^+/+^ male), 8 (hAPPJ20/Bdh1^Thy1−/−^ male), 28 (hAPPJ20/Bdh1^+/+^ female), and 16 (hAPPJ20/Bdh1^Thy1−/−^ female) mice per group. P = 0.020 (male, day 2, t = 2.42, df = 46), p < 0.0001 (male, day 3, t = 4.47, df = 46), p = 0.00063 (male, day 4, t = 3.67, df = 46), p = 0.013 (hAPPJ20 female, day 2, t = 2.61, df = 42), p < 0.0001 (hAPPJ20 female, day 3, t = 4.55, df = 42), p = 0.00050 (hAPPJ20 female, day 4, t = 3.78, df = 42), p = 0.00034 (male, day 5, t = 3.87, df = 46), p < 0.0001 (male, day 13, t = 4.38, df = 46), p = 0.0016 (hAPPJ20 female, day 5, t = 3.38, df = 42), and p = 0.0003 (hAPPJ20 female, day 13, t = 3.92, df = 39) by unpaired two-tailed t-test assuming Gaussian distribution. Data are means ± SD. Mice were given tamoxifen at four months old to knockout *Bdh1* in neurons.

## Discussion

While neurons are hypothesized to rely mainly on astrocyte-derived lactate (*40*) or direct glucose metabolism for energy (*41, 42*), we show that neurons can and must metabolize ketones for normal cellular bioenergetic functions, memory and survival, including under glucose-abundant conditions. In human neurons, physiologic levels of βHB are required for maximum acetyl-CoA generation, mitochondrial oxidation, and respiratory reserve. *In vivo*, mice with adult-induced knockout of Bdh1 in Thy1-expressing cortical neurons die prematurely and display deficits in memory tasks, phenotypes that are further exacerbated in an Alzheimer’s disease model. As such, these data reveal a previously unappreciated obligate requirement for βHB oxidation to AcAc by Bdh1 in neuronal metabolism to support brain function, slow aging, and protect against neurodegeneration.

### Neurons require Bdh1 for physiologic energy metabolism

The constitutive function of ketone bodies, beyond their roles in fasting or specialized diets, remains understudied outside early development (*9, 10*). Our dual use of mitochondrial assays in human neurons *in vitro*, and cell-specific, adult-onset knockouts and longitudinal lifespan analyses *in vivo*, enabled us to uncover this requirement. Prior studies manipulating Bdh1 or Oxct1 used germline, whole-body, or liver-specific knockouts, which were confounded by developmental effects, systemic or liver disease, or limited to early adulthood (*43-49*). By contrast, our Thy1-driven conditional model preserves normal systemic ketone levels and liver function, allowing us to dissociate βHB supply from its utilization within neurons.

The unclear extent of *in situ* ketogenesis in the brain may have obscured the constitutive requirement for ketone metabolism we observed. The liver generates nearly all circulating ketone bodies (*2*), readily measured via arteriovenous differences as hepatocytes lack Oxct1, the enzyme required for ketone oxidation (*5*). Assuming peripheral ketones are the brain’s primary source, minimal arteriovenous differences in humans undergoing brain surgery (*50*) and in fed rats (*51*) argue against substantial brain ketone utilization in non-starved states. Brain uptake of ketone tracers correlates with circulating levels, which are low under basal conditions (*52*). However, *in situ* ketogenesis by astrocytes, as suggested by *in vitro* (*6, 7*) and model organism studies (*8*), and analogous to the lactate shuttle (*40*), could reconcile these findings. Clarifying the extent and function of brain ketogenesis remains an important direction for future research, with potential as a therapeutic target for neurodegenerative disease.

Recent studies show that acute βHB addition supports neural activity and boosts mitochondrial respiration (*53, 54*) in primary cultures, although the relative contributions from astrocytes versus neurons to βHB metabolism were not investigated. We found that βHB contributes to acetyl-CoA and TCA intermediates in human neurons *in vitro*, in a Bdh1-dependent manner. Loss of βHB metabolism to acetoacetate significantly decreases TCA metabolites, with the extent of reduction dependent on the relative availability of glucose and βHB, highlighting the potential to modulate brain energy via dietary fuels.

Notably, AcAc supplementation increased TCA metabolites and respiration more than did βHB. Since βHB oxidation depends on Bdh1-mediated conversion to AcAc, the stronger effect of AcAc may reflect more efficient metabolism. Alternatively, AcAc may also have oxidation-independent signaling roles that regulate metabolism (*4*). These signaling effects may also contribute to the differing potency of AcAc and βHB, and is an important area for future investigation. Under constitutive conditions, blood concentrations of βHB and AcAc are similar (*2*). For our *in vivo* experiments we selected Bdh1 rather than Oxct1 (which would abrogate both βHB and AcAc oxidation) to moderately reduce overall ketone body oxidation in neurons, based on our observation of sensitive dose-responsive changes in metabolism, and because our *in vitro* data suggested that AcAc compensation for Bdh1 knockdown was incomplete (fig. 3).

### Ketone metabolism in AD pathophysiology

Our findings extend beyond normal physiology to implicate neuronal ketone metabolism in protection against neurodegeneration. While BDH1 mutations have not yet been linked directly to human disease, the gene lies within the 3q29 locus implicated in microdeletion and microduplication neurodevelopmental disorders (*55, 56*). BDH1 expression is also reduced in multiple brain cell types in AD (*57*). Moreover, polymorphisms in BDH1 are associated with progression from mild cognitive impairment to AD (*58*), supporting a potential role in disease susceptibility.

Our finding that BDH1 deficiency in neurons markedly increases the mortality of mutant APP (J20) mice supports that constitutive neuronal ketone metabolism may normally protect against AD pathophysiology. Considerable evidence supports that mutant APP can disrupt energy metabolism in mouse models and in patients (*12, 59*), while supplementation with ketones can improve function in AD mouse models (*60, 61*). As such, ketones may provide an alternate fuel source that bypasses mutant APP-induced energetic deficits. Importantly, we show here that this is not only relevant to high-dose exogenous supplementation of ketones, but that the brain’s intrinsic, constitutive ketone metabolism provides crucial endogenous resilience to AD deficits. Feeding a ketone ester to Alzheimer’s mouse models has been shown to reduce seizure activity (*62, 63*) which is linked to memory deficits in humans with AD (*64*), but also a major cause of death in AD mouse models. Exacerbation of this seizure activity in the absence of constitutive neuronal BHB oxidation could be a mechanism of accelerated mortality in hAPPJ20/Bdh1^Thy1−/−^ mice. It is of keen interest to determine if the same or distinct mechanisms are responsible for βHB oxidation and BDH1 activity in improving health and survival during normal aging and in neurodegeneration models.

Altogether, our findings support the possibility that enabling or increasing constitutive ketone body metabolism in neurons could be protective against age-associated cognitive decline and AD. While various early clinical trials are investigating exogenously provided ketone bodies in AD (*12, 65, 66*), targeting endogenous ketogenic pathways may prove equally or more efficacious. Our findings illustrate the need to better understand glial ketogenesis, constitutive neuronal ketone oxidation, and the distinct roles of βHB, AcAc, and other non-glucose fuels in supporting normal neuronal function in aging and neurodegeneration.

## Acknowledgments

We thank Johanna ten Hoeve-Scott, Thomas G. Graeber, and the UCLA Metabolomics Center for their assistance with the metabolomics studies and data processing. We also thank Francoise Chanut for helping edit the manuscript, and Tami Tolpa for designing the figure schemas. This work was supported by funding from the following sources: National Institute on Aging & National Institutes of Health grant numbers R01AG067333 (JCN), R01 AG065428 (KN), RF1 AG064170 (KN), and T32AG000266 (MN); Longevity Impetus (JCN);

UCSF Bakar Aging Research Institute (KN); Hillblom Center & Bakar Aging Research Institute Graduate Fellowship (JY); UCSF Nutrition Obesity Research Center (KN); Hevolution Foundation (JCN); and Buck Institute institutional funds (JCN).

## Author contributions

J.Y. and M.N. contributed to conceptualization, investigation, visualization, funding acquisition, and manuscript preparation; J.X.M. contributed to methodology, investigation, and visualization; T.Y.G. contributed to investigation and project administration; T.R.M. and D.P.K. contributed to methodology; and K.N. and J.C.N. contributed to conceptualization, funding acquisition, supervision, and manuscript preparation.

## Data Availability, Materials & Correspondence

All data are available in the manuscript or the supplementary materials. All data, code, and materials used in the analysis will be made available to any researcher for purposes of reproducing or extending the analyses.

Correspondence and material requests should be addressed to Ken Nakamura and John C Newman.

## References

1. O. E. Owen et al., Brain metabolism during fasting. J Clin Invest 46, 1589–1595 (1967).

2. G. F. Cahill, Jr., Fuel metabolism in starvation. Annu Rev Nutr 26, 1–22 (2006).

3. G. A. Brooks, The Science and Translation of Lactate Shuttle Theory. Cell Metab 27, 757–785 (2018).

4. A. B. Nelson, E. D. Queathem, P. Puchalska, P. A. Crawford, Metabolic Messengers: ketone bodies. Nat Metab 5, 2062–2074 (2023).

5. J. C. Newman, E. Verdin, β-Hydroxybutyrate: A Signaling Metabolite. Annu Rev Nutr 37, 51–76 (2017).

6. N. Auestad, R. A. Korsak, J. W. Morrow, J. Edmond, Fatty acid oxidation and ketogenesis by astrocytes in primary culture. J Neurochem 56, 1376–1386 (1991).

7. C. Blazquez, C. Sanchez, G. Velasco, M. Guzman, Role of carnitine palmitoyltransferase I in the control of ketogenesis in primary cultures of rat astrocytes. J Neurochem 71, 1597–1606 (1998).

8. B. Silva et al., Glia fuel neurons with locally synthesized ketone bodies to sustain memory under starvation. Nat Metab 4, 213–224 (2022).

9. D. G. Cotter, D. A. d’Avignon, A. E. Wentz, M. L. Weber, P. A. Crawford, Obligate role for ketone body oxidation in neonatal metabolic homeostasis. J Biol Chem 286, 6902–6910 (2011).

10. A. Nehlig, Brain uptake and metabolism of ketone bodies in animal models. Prostaglandins Leukot Essent Fatty Acids 70, 265–275 (2004).

11. S. Camandola, M. P. Mattson, Brain metabolism in health, aging, and neurodegeneration. Embo j 36, 1474–1492 (2017).

12. S. C. Cunnane et al., Brain energy rescue: an emerging therapeutic concept for neurodegenerative disorders of ageing. Nat Rev Drug Discov 19, 609–633 (2020).

13. L. Mosconi et al., Hippocampal hypometabolism predicts cognitive decline from normal aging. Neurobiol Aging 29, 676–692 (2008).

14. E. M. Reiman et al., Preclinical evidence of Alzheimer’s disease in persons homozygous for the epsilon 4 allele for apolipoprotein E. N Engl J Med 334, 752–758 (1996).

15. J. Yao et al., Mitochondrial bioenergetic deficit precedes Alzheimer’s pathology in female mouse model of Alzheimer’s disease. Proc Natl Acad Sci U S A 106, 14670–14675 (2009).

16. L. Mosconi et al., Hypometabolism and altered cerebrospinal fluid markers in normal apolipoprotein E E4 carriers with subjective memory complaints. Biol Psychiatry 63, 609–618 (2008).

17. E. M. Reiman et al., Functional brain abnormalities in young adults at genetic risk for late-onset Alzheimer’s dementia. Proc Natl Acad Sci U S A 101, 284–289 (2004).

18. D. Ebert, R. G. Haller, M. E. Walton, Energy contribution of octanoate to intact rat brain metabolism measured by 13C nuclear magnetic resonance spectroscopy. J Neurosci 23, 5928–5935 (2003).

19. S. Hoyer, Abnormalities of glucose metabolism in Alzheimer’s disease. Ann N Y Acad Sci 640, 53–58 (1991).

20. E. Croteau et al., Ketogenic Medium Chain Triglycerides Increase Brain Energy Metabolism in Alzheimer’s Disease. J Alzheimers Dis 64, 551–561 (2018).

21. T. R. Matsuura, P. Puchalska, P. A. Crawford, D. P. Kelly, Ketones and the Heart: Metabolic Principles and Therapeutic Implications. Circ Res 132, 882–898 (2023).

22. J. L. Horton et al., The failing heart utilizes 3-hydroxybutyrate as a metabolic stress defense. JCI Insight 4, (2019).

23. L. Rong et al., Effects of ketogenic diet on cognitive function of patients with Alzheimer’s disease: a systematic review and meta-analysis. J Nutr Health Aging 28, 100306 (2024).

24. M. G. Grammatikopoulou et al., To Keto or Not to Keto? A Systematic Review of Randomized Controlled Trials Assessing the Effects of Ketogenic Therapy on Alzheimer Disease. Adv Nutr 11, 1583–1602 (2020).

25. C. Wang et al., Scalable Production of iPSC-Derived Human Neurons to Identify Tau-Lowering Compounds by High-Content Screening. Stem Cell Reports 9, 1221–1233 (2017).

26. Y. Zhang et al., Rapid single-step induction of functional neurons from human pluripotent stem cells. Neuron 78, 785–798 (2013).

27. R. Tian et al., CRISPR Interference-Based Platform for Multimodal Genetic Screens in Human iPSC-Derived Neurons. Neuron 104, 239–255 e212 (2019).

28. H. Langemann et al., Extracellular levels of glucose and lactate measured by quantitative microdialysis in the human brain. Neurol Res 23, 531–536 (2001).

29. J. W. Pan et al., [2,4-13 C2]-beta-Hydroxybutyrate metabolism in human brain. J Cereb Blood Flow Metab 22, 890–898 (2002).

30. G. A. Dienel, Brain lactate metabolism: the discoveries and the controversies. J Cereb Blood Flow Metab 32, 1107–1138 (2012).

31. E. C. McNay, T. M. Fries, P. E. Gold, Decreases in rat extracellular hippocampal glucose concentration associated with cognitive demand during a spatial task. Proceedings of the National Academy of Sciences 97, 2881–2885 (2000).

32. A. Rex, B. Bert, H. Fink, J.-P. Voigt, Stimulus-dependent changes of extracellular glucose in the rat hippocampus determined by in vivo microdialysis. Physiol Behav 98, 467–473 (2009).

33. D. Malinowska, M. Żendzian-Piotrowska, Ketogenic Diet: A Review of Composition Diversity, Mechanism of Action and Clinical Application. J Nutr Metab 2024, 6666171 (2024).

34. P. Tognini et al., Distinct Circadian Signatures in Liver and Gut Clocks Revealed by Ketogenic Diet. Cell Metab 26, 523-538.e525 (2017).

35. P. Young et al., Single-neuron labeling with inducible Cre-mediated knockout in transgenic mice. Nat Neurosci 11, 721–728 (2008).

36. A. Lee et al., Abeta42 oligomers trigger synaptic loss through CAMKK2-AMPK-dependent effectors coordinating mitochondrial fission and mitophagy. Nat Commun 13, 4444 (2022).

37. S. J. Beck et al., Deregulation of mitochondrial F1FO-ATP synthase via OSCP in Alzheimer’s disease. Nat Commun 7, 11483 (2016).

38. L. Chen et al., Studies on APP metabolism related to age-associated mitochondrial dysfunction in APP/PS1 transgenic mice. Aging (Albany NY) 11, 10242–10251 (2019).

39. L. Mucke et al., High-level neuronal expression of abeta 1-42 in wild-type human amyloid protein precursor transgenic mice: synaptotoxicity without plaque formation. J Neurosci 20, 4050–4058 (2000).

40. M. Belanger, I. Allaman, P. J. Magistretti, Brain energy metabolism: focus on astrocyte-neuron metabolic cooperation. Cell Metab 14, 724–738 (2011).

41. C. M. Diaz-Garcia et al., Neuronal Stimulation Triggers Neuronal Glycolysis and Not Lactate Uptake. Cell Metab 26, 361–374 e364 (2017).

42. H. Li et al., Neurons require glucose uptake and glycolysis in vivo. Cell Rep 42, 112335 (2023).

43. Y. Arima et al., Murine neonatal ketogenesis preserves mitochondrial energetics by preventing protein hyperacetylation. Nat Metab 3, 196–210 (2021).

44. S. Asif et al., Hmgcs2-mediated ketogenesis modulates high-fat diet-induced hepatosteatosis. Mol Metab 61, 101494 (2022).

45. H. Otsuka et al., Deficiency of 3-hydroxybutyrate dehydrogenase (BDH1) in mice causes low ketone body levels and fatty liver during fasting. J Inherit Metab Dis 43, 960–968 (2020).

46. D. B. Stagg et al., Diminished ketone interconversion, hepatic TCA cycle flux, and glucose production in D-beta-hydroxybutyrate dehydrogenase hepatocyte-deficient mice. Mol Metab 53, 101269 (2021).

47. D. G. Cotter, R. C. Schugar, A. E. Wentz, D. A. d’Avignon, P. A. Crawford, Successful adaptation to ketosis by mice with tissue-specific deficiency of ketone body oxidation. Am J Physiol Endocrinol Metab 304, E363–374 (2013).

48. J. Enders et al., Ketolysis is required for the proper development and function of the somatosensory nervous system. Exp Neurol 365, 114428 (2023).

49. I. Tomita et al., Ketone bodies: A double-edged sword for mammalian life span. Aging Cell 22, e13833 (2023).

50. Y. Wang et al., Comprehensive characterization of metabolic consumption and production by the human brain. Neuron 113, 1708-1722.e1705 (2025).

51. N. B. Ruderman, P. S. Ross, M. Berger, M. N. Goodman, Regulation of glucose and ketone-body metabolism in brain of anaesthetized rats. Biochem J 138, 1–10 (1974).

52. S. C. Cunnane et al., Can ketones compensate for deteriorating brain glucose uptake during aging? Implications for the risk and treatment of Alzheimer’s disease. Ann N Y Acad Sci 1367, 12–20 (2016).

53. K. Bredvik, C. Liu, T. A. Ryan, Characterization of β-Hydroxybutyrate as a Cell Autonomous Fuel for Active Excitatory and Inhibitory Neurons: β-Hydroxybutyrate as a Fuel for Active Neurons. bioRxiv, (2024).

54. I. Llorente-Folch, H. Düssmann, O. Watters, N. M. C. Connolly, J. H. M. Prehn, Ketone body β-hydroxybutyrate (BHB) preserves mitochondrial bioenergetics. Sci Rep 13, 19664 (2023).

55. A. A. Kashevarova et al., Delineation of the Genetic Architecture and Clinical Polymorphism of 3q29 Duplication Syndrome: A Review of the Literature and a Report of Two Novel Patients With Single-Gene BDH1 Duplications. Mol Genet Genomic Med 13, e70047 (2025).

56. M. Massier et al., 3q29 duplications: A cohort of 46 patients and a literature review. Am J Med Genet A 194, e63531 (2024).

57. H. Mathys et al., Single-cell atlas reveals correlates of high cognitive function, dementia, and resilience to Alzheimer’s disease pathology. Cell 186, 4365–4385 e4327 (2023).

58. E. Lee et al., Single-nucleotide polymorphisms are associated with cognitive decline at Alzheimer’s disease conversion within mild cognitive impairment patients. Alzheimers Dement (Amst) 8, 86–95 (2017).

59. E. Tönnies, E. Trushina, Oxidative Stress, Synaptic Dysfunction, and Alzheimer’s Disease. J Alzheimers Dis 57, 1105–1121 (2017).

60. Y. Kashiwaya et al., A ketone ester diet exhibits anxiolytic and cognition-sparing properties, and lessens amyloid and tau pathologies in a mouse model of Alzheimer’s disease. Neurobiol Aging 34, 1530–1539 (2013).

61. J. Di Lucente et al., Ketogenic diet and BHB rescue the fall of long-term potentiation in an Alzheimer’s mouse model and stimulates synaptic plasticity pathway enzymes. Commun Biol 7, 195 (2024).

62. J. C. Newman et al., Small-molecule ketone esters treat brain network abnormalities in an Alzheimer’s disease mouse model. bioRxiv, 2022.2009.2024.509337 (2022).

63. A. Cheng et al., SIRT3 Haploinsufficiency Aggravates Loss of GABAergic Interneurons and Neuronal Network Hyperexcitability in an Alzheimer’s Disease Model. J Neurosci 40, 694–709 (2020).

64. J. J. Palop, L. Mucke, Network abnormalities and interneuron dysfunction in Alzheimer disease. Nat Rev Neurosci 17, 777–792 (2016).

65. P. Krolak-Salmon et al., Efficacy and Safety of Exogenous Ketones in People with Mild Neurocognitive Disorder and Alzheimer’s Disease: A Systematic Literature Review. Nutr Rev 83, e1034–e1048 (2025).

66. É. Myette-Côté, A. Soto-Mota, S. C. Cunnane, Ketones: potential to achieve brain energy rescue and sustain cognitive health during ageing. Br J Nutr 128, 407–423 (2022).

