## Supplementary material for "Requirement for oxidation of neuronal ketone bodies in aging and neurodegeneration": Methods and Extended Data

Neuronal differentiation and maturation

These studies were conducted in human WTC11 induced pluripotent stem cells (iPSCs) expressing doxycycline-inducible neurogenin 2 (NGN2) and inducible CRISPRi (dCas9-KRAB) machinery, obtained from Martin Kampmann (UCSF). iPSCs were cultured in StemFlex media (Gibco A3349401) on Matrigel-coated (1:120 in DMEM/F12; Corning 354277, Gibco 11320033) culture flasks. Differentiation of the iPSCs was divided into 2 parts: pre-differentiation and maturation. Pre-differentiation occurred over 3d of culture in pre-differentiation media; maturation occurred over 18-22d in neuronal differentiation media. Pre-differentiation media was based in Knockout DMEM/F12 (Gibco 12660012), with 1X NEAA (Gibco 11140050), 1X N2 supplement (Gibco 17502001), 10 ng/mL NT-3 (StemCell 78074.1), 10 ng/mL BDNF (StemCell 78005.1), and 1 μg/mL of laminin (Gibco 23017015). On the first day of pre-differentiation, iPSCs were passaged onto Matrigel-coated flasks with pre-differentiation media containing doxycycline (2 μg/mL) and ROCK inhibitor (ROCKi; 10 μM). Media was refreshed on the third day of pre-differentiation to remove ROCKi.

On the fourth day, pre-differentiated neurons were passaged into 6- or 96-well plates with neuronal differentiation media and doxycycline (2 μg/mL), at densities of 500,000 or 10,000-12,000 cells/well, respectively. Plates were pretreated for 2d with 1X poly-L-ornithine (Sigma-Aldrich P4957) and coated for 1d with 5 μg/mL laminin. Neuronal differentiation media was based in 1:1 DMEM/F12 (Gibco 11320033) and Neurobasal A media (Gibco 10888022), with 1X NEAA, 0.5X GlutaMAX (Gibco 35050061), 0.5X N2 supplement, 0.5X B27 supplement (Gibco 17504001), 10 ng/mL NT-3, 10 ng/mL BDNF, and 1 μg/mL of laminin. After 2d, media was refreshed to remove doxycycline and then refreshed every 7d to feed the neurons.

Minimal base media for metabolic restriction

All metabolic conditions were created in an absolute minimal media. The media was based in Neurobasal A -glucose -pyruvate (Gibco A2477501), with 1X NEAA (Gibco 11140050), 0.375X GlutaMAX (Gibco 35050061), 0.5X N2 supplement (Gibco 17502001), 0.5X B27 supplement (Gibco 17504001), 10 ng/mL NT-3 (StemCell 78074.1), 10 ng/mL BDNF (StemCell 78005.1), and 1 μg/mL laminin (Gibco 23017015).

Timescale targeted metabolomics in ketogenic conditions

WTC11 human neurons were differentiated in 6-well plates, matured for 20d, then incubated in metabolic media prior to metabolite extraction. On day 20, media was changed to one of the following metabolic conditions, with 4 replicates each:

[1] Physiologic – 1.5mM glucose (Sigma-Aldrich G6152), 0.2mM pyruvate (Sigma-Aldrich P5280), 0.2mM BHB (Sigma-Aldrich H6501), 1mM lactate (Sigma-Aldrich L7022)

[2] Ketogenic – 0.5mM glucose, 0.2mM pyruvate, 3mM BHB

[3] Extreme ketogenic – 10mM BHB only

Conditions were run in duplicate with [U-13C] versions of either glucose (Cambridge Isotope Labs CLM-1396) or BHB (Cambridge Isotope Labs CLM-3853). All conditions were formulated in previously-described minimal base media. For Acetyl-CoA measurements, an additional condition of 1.5mM glucose, 0.2mM pyruvate, 10mM BHB, and 1mM lactate was assayed. After 4, 24, or 48h of metabolic media incubation, cellular metabolites were extracted via ammonium acetate wash (150mM, pH 7.4) at 4C and incubation with 80% methanol for 30min at -80C, followed by centrifugation at 14,000 rpm for 10min at 4C. Metabolite supernatants were dried in a Labconco CentriVap prior to quantification at the UCLA Metabolomics Core.

Fatty acid conjugation

For all fatty acid-based metabolic medias, fatty acids were first resuspended in 100% ethanol and slowly added to 10% fatty acid-free BSA (Roche 03117057001) in minimal base media at 37C, for a final concentration of 2.5mM (palmitate) or 5mM (oleate, stearate). Solutions were vortexed vigorously and incubated for >30min at 37C. Control conditions with 10% BSA, ethanol, or both in base media were also included.

Survival assays

Human neurons were differentiated in 96-well plates, matured for 18 days, then incubated for 48 or 72h in metabolic media prior to assaying for cell death. On day 18, media was changed to metabolic conditions formulated in minimal base media, with increasing doses of BHB (Sigma-Aldrich H6501), AcAc (Sigma-Aldrich A8509), Acetone (Fisher Scientific A18-4), palmitate (Sigma-Aldrich P0500), oleate (Sigma-Aldrich O1008), or stearate (Sigma-Aldrich S4751).

After media incubation, cells were incubated with Calcein (Invitrogen C3099) and Hoechst (Invitrogen H3570) at 37C for 15min prior to imaging on the Thermo Scientific CellInsight CX7 microscope. Images were analyzed in CellProfiler to determine survival via colocalization of Calcein and Hoechst. Counts were normalized to average survival in the lowest concentration (at 48h, for fuel assays; in NTG1, for BDH1 assays) per experimental replicate and combined to generate dose curves.

Targeted metabolomics

Human neurons were differentiated in 6-well plates, matured for 18d, then incubated for 3d in metabolic media prior to metabolite extraction. On day 18, media was changed to one of the following metabolic conditions, with 4-6 replicates each:

[1] Basal (unrestricted) – 21.25mM [U-13C]-glucose and 2.5mM pyruvate
[2] Physiologic – 1.5mM glc and 0.2mM BHB, with [U-13C]-glc or -BHB
[3] Physiologic variant – 1.5mM glc and 0.2mM AcAc, with [U-13C]-glc or -AcAc
[4] Ketone reliance – 3mM or 8mM [U-13C]-BHB, -AcAc (Supelco 77988), or -acetone (Cambridge Isotope Labs CLM-1334)
[5] Fatty acid reliance – 50µM [U-13C]-palmitate (Sigma-Aldrich 605573), -oleate (Cambridge Isotope Labs CLM-460), or -stearate (Sigma-Aldrich 605581)
[6] Competition – 3mM glc, BHB, and AcAc; run in triplicate with [U-13C] per fuel

All conditions were formulated in previously-described minimal base media; basal condition contained extra GlutaMAX to a final concentration of 1.125X. After 72h of media incubation, cellular metabolites were extracted and quantified at the UCLA Metabolomics Core. Values were normalized to corresponding physiologic condition per tracer run.

Seahorse

WTC11 human neurons were differentiated in Seahorse XFe96/XF Pro Cell Culture Microplates (Agilent 103792-100) as previously described, matured for 18d, then incubated for 3d in metabolic media prior to Seahorse assay. On day 18, media was changed to one of the following metabolic conditions:

[1] Physiologic – 1.5mM glc and 0.2mM BHB
[2] BHB variants – 1.5mM glc and varying doses of BHB
[3] Glucose variants – 0.2mM BHB and varying doses of glucose
[4] Ketone reliance – 8mM BHB, AcAc, or acetone
[5] Fatty acid reliance – 50µM palmitate, oleate, or stearate

After 72h of metabolic media incubation, all conditions were remade in minimal media, but with Seahorse XF DMEM (Agilent 103575-100) as the base media instead of Neurobasal A. Plates were incubated in CO2-free incubator at 37C for 1-2h prior to loading into Seahorse XF96 Extracellular Flux Analyzer. Seahorse sensor cartridges (Agilent 103792-100) were calibrated per Agilent instructions. Oxygen consumption rate (OCR) was measured at baseline and again after sequential injection of DMSO (Sigma-Aldrich D2438), FCCP (carbonyl cyanide-4-(trifluoromethoxy)phenylhydrazone, Sigma-Aldrich C2920), rotenone (Sigma-Aldrich R8875), and DMSO. OCR signals were normalized to average baseline reading per replicate.

Generation of BDH1 knockdown cell line

The dual-guide RNA expression plasmid pJR103 (Addgene plasmid #187242) was used as the lentiviral transfer vector. The following protospacer sequences were cloned into pJR103: for the non-targeting control (NTG1), 5′-GGGCTAAGGGGCCGTGTACT-3′ and 5′-GGGACGAAGCAGTCCGAGCG-3′; for BDH1 knockdown (BDH1-KD), 5′-GCGCAGGAGTGCTGGTGGAG-3′ and 5′-GGCGTGTAGAAGCGTCCGGG-3′.

Lentivirus was produced by transient transfection of HEK293T cells (ATCC CRL-11268). Cells were seeded to reach 80–95% confluence after 24h. For each well of a 6-well plate, 1 μg of pJR103 transfer plasmid and 1 μg of third-generation packaging plasmid mix [equal ratio of pRSV-Rev (Adgene plasmid #12253), pMDLg (Adgene plasmid #12251), pCMV-VSV-G (Adgene plasmid #8454)] were diluted in 200 μL OPTI-MEM (Thermo Fisher Scientific 31985070). TransIT-Lenti transfection reagent (Mirus Bio MIR 6600) was added at 6 μL per well and incubated at RT for 10min before being added dropwise to HEK293T cells.

Viral supernatants were harvested at 48h and 72h post-transfection, filtered through a 0.45 μm PVDF syringe filter (Millipore Sigma SLHV033RB), and mixed with one-quarter volume of Lentivirus Precipitation Solution (Alstem VC125). The mixture was incubated at 4 °C for 24h, followed by centrifugation at 1,500 × g for 30min at 4 °C. The supernatant was carefully aspirated, and the pellet centrifuged again for 5min at 1,500 × g to remove residual liquid. The viral pellet was resuspended in 100 μL sterile PBS (UCSF media core).

For transduction, the WTC11 iPSCs were infected with concentrated lentivirus at the time of passaging. 48h post-infection, cells were passaged into fresh medium containing 1 μg/mL puromycin (Sigma Aldrich P9620) and selected for two passages until greater than 95% of the population was marker-positive, as assessed by flow cytometry. Cells were then passaged once in puromycin-free medium to allow recovery prior to subsequent differentiation experiments.

Confirmation of BDH1 knockdown

Total RNA was isolated from cells collected at multiple timepoints following lentiviral transduction. RNA was extracted using the RNeasy Plus Mini Kit (QIAGEN 74134) and reverse-transcribed using the High-Capacity RNA-to-cDNA™ Kit (Thermo Fisher Scientific 4387406) in a 20 μL reaction following the manufacturer's protocol. Quantitative PCR was performed using TaqMan™ Gene Expression Assays (Thermo Fisher Scientific 4444557) on a QuantStudio 5 Real-Time PCR System (Applied Biosystems). Reactions were carried out in technical triplicate. The TaqMan™ assay for BDH1 (Hs00366297_m1) was used to quantify target gene expression, with ACTB (β-actin; Hs99999903_m1) as the endogenous control. Relative BDH1 expression was calculated using the comparative Ct (ΔΔCt) method, normalizing to ACTB. Expression levels were compared to those of cells transduced with non-targeting guide (NTG1) controls at day 0 to determine knockdown efficiency.

Mouse strains, housing, and husbandry

All mice were maintained in accordance with the guidelines set forth by the National Institutes of Health, and all experimental protocols were approved by the Buck Institute Institutional Animal Care and Use Committee (IACUC). The Buck Institute is accredited by the Association for Assessment and Accreditation of Laboratory Animal Care (AAALAC). Mice were maintained in a specific pathogen-free barrier facility under a 6:00 am to 6:00 pm light cycle. The facility maintains a temperature of 68-72°F and an ambient humidity level exceeding 30%. All mice were maintained in groups of up to five mice per cage, unless otherwise specified in procedures described below. The cages were furnished with wooden bedding, a nestlet, and a house, and were changed every two weeks. The diet was provided in a recess of the metal wire lid situated at the upper portion of the cage and changed every two weeks, and the wire lid was changed with the diet. Water was provided in a bottle. Bdh1$^{fl/fl}$ mice were previously published (*22*). Thy1-CreERT2 (SLICK-H) mice (Strain #:012708) (*35*) and APP/J20 mice (Strain #034836-JAX) (*39*) were obtained from the Jackson Laboratory. The age of mice at the time experiments were undertaken is presented in the figures. Knockout of *Bdh1* was induced by intraperitoneal tamoxifen injections (25 mg/ml corn oil, 2.5 mg per 25 g of body weight) on five consecutive days. The lifespan cohorts began with four months old mice after tamoxifen injections and followed them until death; the healthspan involved behavioral testing at the described ages.

<u>Mouse diets and feeding</u>

Unless otherwise indicated, mice were fed the Buck Institute standard vivarium chow with 18% protein, 6% fat, and 44% carbohydrates (2918; Teklad). The customized diets from Envigo contained the following macronutrient content per calorie: Control (CD) with 10% protein, 13% fat, and 77% carbohydrate (TD.150345); KD with 10% protein and 90% fat (TD.160153). Both diets had similar micronutrient content, fiber, and preservatives on a per-calorie basis, had similar fat sources, and were always provided ad libitum.

<u>Immunofluorescence staining</u>

The brains were fixed with 4% PFA and subsequently sectioned at a thickness of 30 μm using a microtome. The brain sections were blocked with 10% normal donkey serum and 0.5% triton-X. The sections were incubated with primary antibodies, mouse anti-NeuN (1:250) (ab104224; Abcam), or mouse anti-GFAP (1:250) (G3893; Sigma), and rabbit anti-BDH1 (1:250) (15417-1-AP; Proteintech), overnight at 4C. Subsequently, the sections were incubated with secondary antibodies, anti-mouse Alexa Fluor 488-conjugated (1:500) (A-21202; Invitrogen) and anti-rabbit Alexa Fluor 555-conjugated (1:500) (A-31572; Invitrogen) and mounted with a solution containing DAPI. The slides were photographed using a Zeiss LSM 700, and the resulting images were subsequently quantified using the ImageJ software.

<u>Blood glucose and plasma BHB measurements</u>

Blood glucose was measure via distal tail-snip by an AimStrip Plus Blood Glucose Meter (37231; Germaine Laboratories). Blood was obtained via distal tail-snip (10-40 μL) or, if the animal was being euthanized for tissue collection, by cardiac puncture immediately following euthanasia. Blood was collected into lithium-heparin coated microvettes (16.443.100; Sarstedt), and plasma separated by centrifugation at 1500 x g for 5 min at 4C. Plasma was frozen at −20C until thawed for assay. We confirmed that freeze-thawing had no effect on assay results. We measured BHB using the Stanbio Chemistry Beta-Hydroxybutyrate LiquiColor Reagent (2440-058; Stanbio), run using 3 μL samples in duplicate. We found it critical to subtract baseline absorbance of sample-enzyme mixture prior to adding catalyst, to account for sample hemolysis.

Glucose and insulin tolerance tests

For the glucose tolerance test, mice were fasted overnight and received an intraperitoneal injection of glucose (2 g/kg body weight). For the insulin tolerance test, the mice were fasted for four hours before being injected intraperitoneally with insulin (0.75 U/kg body weight).

Quantitative PCR (qPCR)

RNA was isolated by Quick-RNA MicroPrep Kit (R1051; Zymo Research), cDNA synthesis was carried out with Superscript cDNA Synthesis Kit (1708891; BioRad), and qPCR was performed using iTaq Universal SYBR Green Supermix (1725121; BioRad) in a BioRad CFX96 Real-Time System. Gene expression analyses were normalized to the Rplp0 housekeeping gene. Primers; Rplp0: Fw_GCTTCGTGTTCACCAAGGAGGA, Rv_GTCCTAGACCAGTGTTCTGAGC; Hmgcs2: Fw_TGCTATGCAGCCTACCGCAAGA, Rv_GCCAGGGATTTCTGGACCATCT: Hmgcl: Fw_GCACTTTGCCAAAGCAGGTGAAG, Rv_CGGAAAGCATGTCGATCAGCCT; Bdh1: Fw_ AGGCTGTGACTCTGGATTTGGG, Rv_CTGGATGGTTCTCAGTCGGTCA; Oxct1: Fw_GAGCGACAGTTCCTTTCTGGTG, Rv_TCCCATACCCTGTGCTGGTGTA.

Lifespan endpoints

The endpoints for a healthy lifespan were based on the NIA Interventions Testing Program (https://www.nia.nih.gov/research/dab/interventions-testing-program-itp), with additional rigorous criteria developed in collaboration with the Buck Vivarium. The goal of the euthanasia criteria was to relieve and prevent any suffering without unnecessarily reducing healthy lifespan. Indications for euthanasia involved obvious discomfort, impending death, systemic signs of unwellness, or any condition that was likely a harbinger of impending discomfort or death. Benign conditions of aging, even if abnormal in young mice, did not prompt euthanasia unless they were associated with these criteria or thought likely to lead to them over days or weeks. All mice were weighed weekly and monitored with increasing frequency up to five times a week after 24 months old. Specific indications for euthanasia of aged mice included the following:

Moribund
BCS 2 or less
Poor general health
Poor mobility
Any obvious discomfort
Discoordination, ataxia, or abnormal gait
Progressive weight loss to > 15% below normal body weight
Unexplained weight gain > 15% above prior body weight
Wounds that do not heal after two weeks of treatment, or worsen despite treatment
Development of new wounds while on treatment for other wounds
Full-thickness wounds that expose underlying fascia or muscle, wounds > 2cm in any dimension, or any wound that is obviously causing discomfort
Rectal prolapse, penile prolapse, or paraphimosis
Suspected tumor associated with other signs of systemic unwellness (wound, weight loss, etc.) Suspected tumor > 2cm or ulcerated

Healthspan study

All behavioral testing was performed in consultation with the Buck Mouse Phenotyping Core, incorporating expert recommendations as to type, sequence, and timing of the various behavioral tests. A key factor was their suitability for use with old mice, compatibility for performing multiple tests on the same animals, and suitability for repeat measurements over time. All tests were carried out during normal daytime hours, in normal room light except where noted.

Novel object recognition

The novel object recognition test was conducted in white plastic boxes with dimensions of 40 x 40 x 30 cm (length x width x height). On the first day of the study, the subjects were allowed to acclimate to the testing environment for five minutes. On the second day, the subjects were given a 10-minute period to become familiar with two identical objects (plastic boxes or bottles) and then a 10-minute test in which they were presented with one known and one novel object (one box and one bottle). On the second day of the experiment, a video camera was utilized to document the experimental procedures. The familiarization period and novel object test were conducted six hours apart. The time spent at both the novel and known objects were recorded.

Open field

The open field apparatus (Tru Scan for Open Field Activity Monitoring; Coulbourn Instruments) consisted of Tru Scan test arenas (25 x 25 x 40 cm) (length x width x height). Mice were recorded for 10 min, in dark conditions.

Y-maze

The Y-maze apparatus was constructed of three arms, each measuring 35 x 5 x 20 cm (length x width x height). These arms radiated out from a central platform, which was shaped like a triangle with sides measuring 5 cm. The room was maintained at a low light level. Each mouse was positioned in the center of the maze, oriented toward one of the arms, and permitted to explore the maze freely for eight minutes. The trials were recorded using a video camera.

Elevated plus maze

The elevated plus maze apparatus consists of two platforms joined in a plus shape. Each platform is 74 cm x 5 cm, with a 5 cm square region where they meet. One platform is flat and open. The other platform has 20 cm high walls. The entire apparatus is 62 cm above the floor. A computer tracking system tracked mouse movement through the apparatus, reporting both distance moved and dwell time. The room was kept at low light. Mice were permitted to explore for 5 min.

RotaRod

The RotaRod was performed with a RotaRod apparatus (47650; Ugo Basile), using a two-day protocol. Day 1 involved a training run at fixed 10 rpm. Days 2 involved three data runs. The data runs on day 2 used steady rod acceleration from 5-40 rpm over 5 min. The time was stopped at 5 min or when the mouse fell. Mice were removed if they rotated around the rod twice.

Barnes maze

The Barnes maze consists of a 110 cm diameter round platform, elevated 80 cm from the floor, and 40 holes (each 5 cm in diameter). All holes were opened except for one target hole that led to an escape box located under the platform. Visual cues and white noise were employed to encourage the mice to expeditiously locate the escape box. Testing was conducted in three phases. Adaptation (Day 0): The mice were permitted to explore the platform for a period of 60 seconds. Any mouse that was unable to locate the escape box was guided and allowed to remain in that location for an additional 60 seconds. Learning Acquisition (Days 1–4): Over the course of four consecutive days, the mice were trained to locate the target and enter the escape box within 180 seconds. Each day, the mice underwent two trials. If they did not locate the target or escape box within 180 seconds, their primary latency was recorded as 180 seconds. Probe (Days 5 and 13): On Days 5 and 13, a single 90-second trial was conducted to measure primary latency. If the mice did not locate the target within 90 seconds, their latency was recorded as 90 seconds.

Quantification and statistical analysis

*In vitro* data are presented as the mean ± standard error (SEM) and *in vivo* data are presented as mean ± standard deviation (SD). Unless otherwise specified in the figure legend, metabolomics and Seahorse datasets were analyzed by Brown-Forsythe and Welch one-way ANOVA with Dunnett's T3 multiple comparisons to compare conditions, or 2-way ANOVA with Dunnett's multiple comparisons for changes over time. Survival assays were analyzed by unpaired two-tailed t-test with Welch's correction. For rodent behavioral tests, p-values were calculated by unpaired two-tailed t-test under the assumption of Gaussian distribution. For lifespan experiments, Log-rank (Mantel-Cox) and Gehan-Breslow-Wilcoxon tests were used to calculate p-values. Statistical calculations were conducted using GraphPad Prism 10 (GraphPad Software).

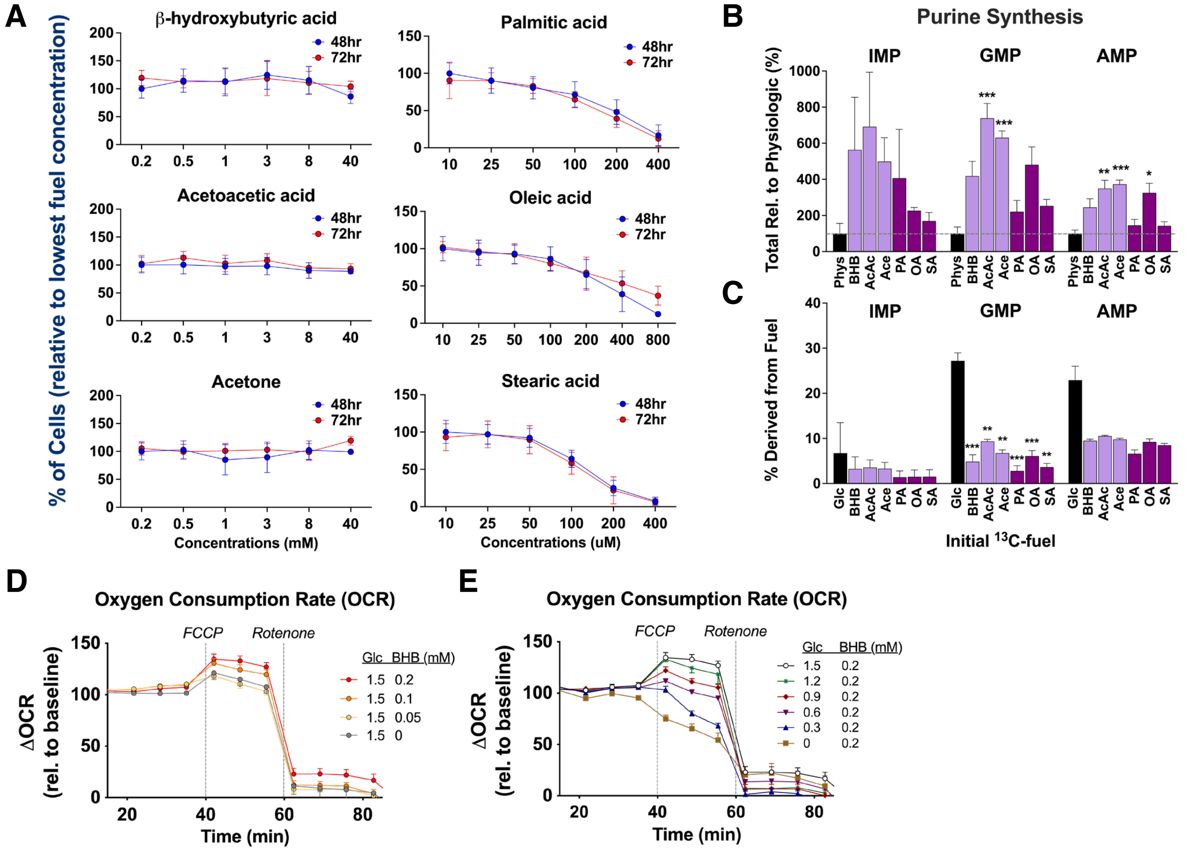


**Extended Data Fig. 1.**

(A) Survival curves of 21d human neurons incubated in lipid-based conditions for 48 or 72h, normalized to lowest fuel concentration at 48h. Conditions include metabolic reliance on ketones (βHB (N=5-11 samples/condition/timepoint), AcAc (5-12, except 3 at 40mM), acetone (5-11, except 2 at 40mM)) or fatty acids (palmitic (10-12), oleic (4-12), stearic acid (7-10)). Data are means ± SD, from two experimental replicates. (B) Total metabolite change relative to physiologic, and (C) percent derivation (from $^{13}$C-fuel) of purine-related species after 3d media incubation. Inosine monophosphate: IMP; guanosine monophosphate: GMP; adenosine monophosphate: AMP. N=3-6 samples/condition. Data are means ± SEM. (D) OCR changes in neurons incubated 3d with incremental changes in βHB or (E) glucose availability. N=3-10 samples/condition. Data are means ± SEM. *$p < 0.05$, **$p < 0.01$, and ***$p < 0.001$ relative to physiologic condition by Brown-Forsythe and Welch one-way ANOVA with Dunnett's T3 multiple comparisons (B-C).

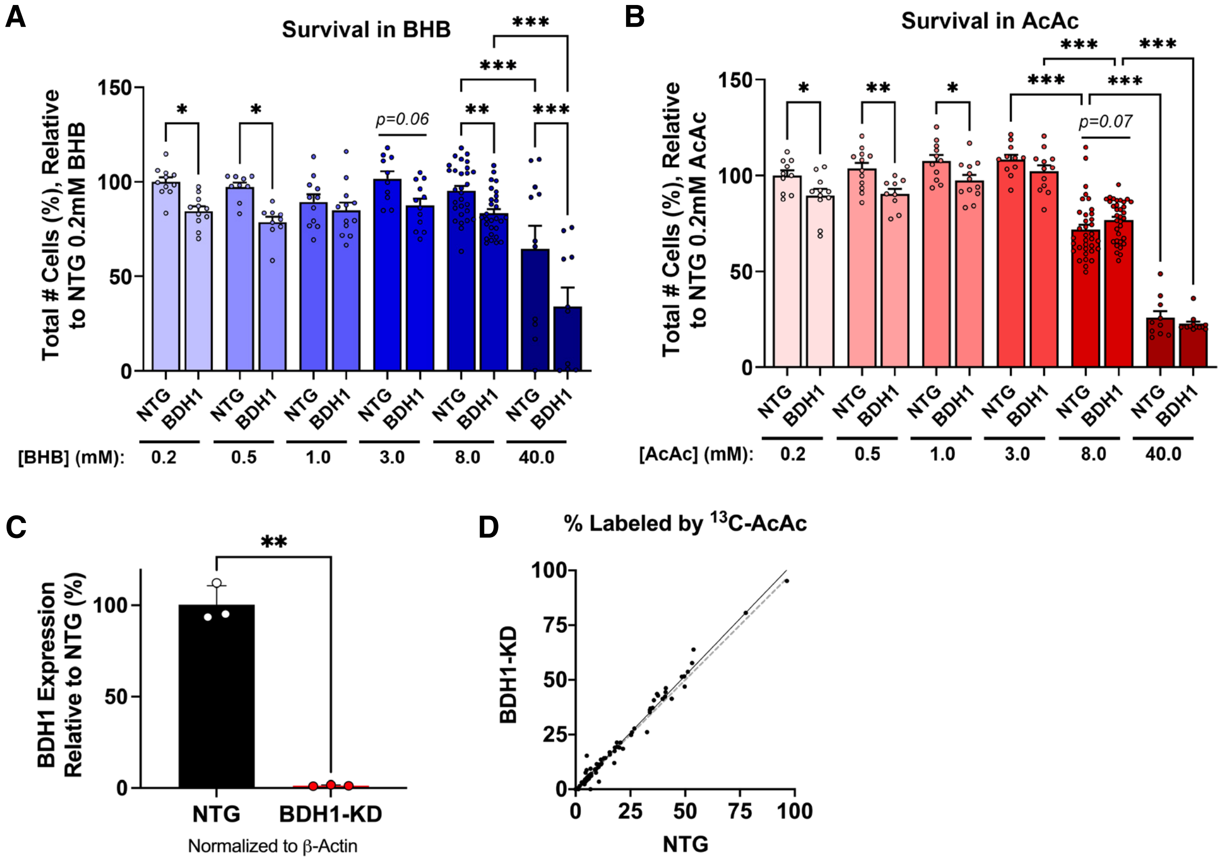


### Extended Data Fig. 2.

Survival curves of 21d NTG or BDH1 neurons incubated for 72hr under (A) βHB-based or (B) AcAc-based metabolic conditions, normalized to NTG survival in 0.2mM ketone per experimental repetition. N=9-35 samples/condition, from two experimental replicates; data are means ± SEM. (C) Validation of *Bdh1* knockdown in BDH1 vs. NTG neurons for targeted metabolomics, normalized to β-Actin, confirming 99% knockdown. N=3 samples/genotype; means ± SD. (D) Regression plot of the BDH1 vs. NTG neuronal metabolome during metabolic reliance on AcAc. *Points:* metabolites; *values:* percent of metabolite containing carbons from $^{13}$C-AcAc. N=115 metabolites; y=1.04x-0.14, $R^2$=0.99. *$p < 0.05$, **$p < 0.01$, and ***$p < 0.001$ by 2-way ANOVA with Tukey's multiple comparisons, select comparisons shown (A-B); or unpaired t-test with Welch's correction (C).

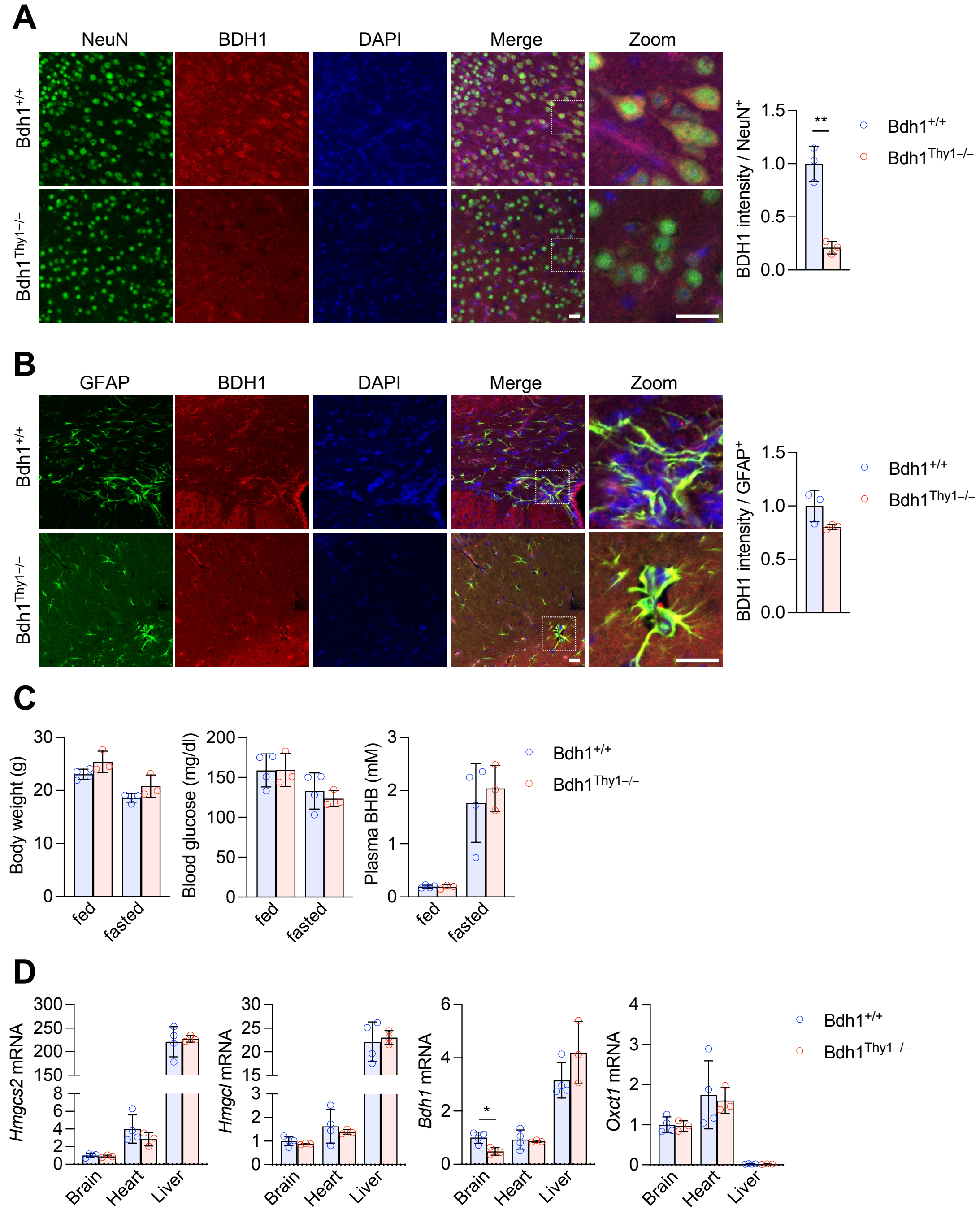


**Extended Data Fig. 3.**

(A and B) Immunostaining of neurons (NeuN) (A) or astrocytes (GFAP) (B), BDH1, and nuclei (DAPI) in brain cortex sections of two-month-old mice harvested after 24 hours fasting and seven days after five consecutive days of tamoxifen injection. BDH1 fluorescent intensity was relatively quantified in cells (N=30 per mouse) that were positive for NeuN (A) or GFAP (B). Scale bar, 25 μm. (A and B) N=3 ($Bdh1^{+/+}$ male) and 3 ($Bdh1^{Thy1-/-}$ male) mice per group. P = 0.0014 (NeuN, t = 7.85, df = 4) by unpaired two-tailed t-test assuming Gaussian distribution. Data are means ± SD. (C) Body weight, blood glucose, and plasma βHB levels before and after 24 hours fasting. (D) qPCR of genes involved in ketone metabolism after 24 hours fasting. (C and D) N=4 ($Bdh1^{+/+}$ male) and 3 ($Bdh1^{Thy1-/-}$ male) mice per group. P = 0.0015 (Bdh1, brain, t = 3.67, df = 5) by unpaired two-tailed t-test assuming Gaussian distribution. Data are means ± SD.

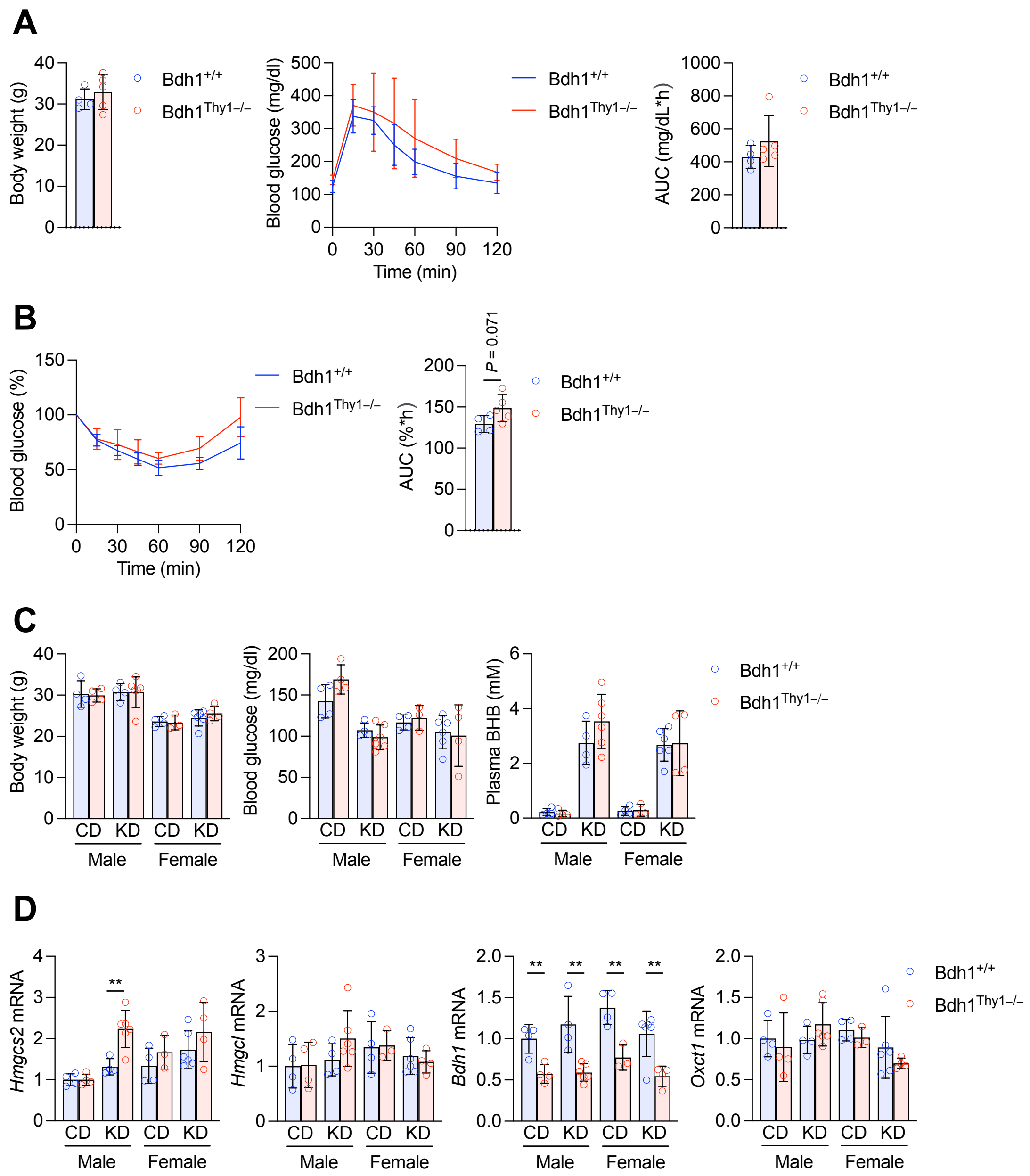


**Extended Data Fig. 4.**

(A) Body weight, glucose tolerance test and the area under the curve (AUC) of nine-month-old mice measured 15 days after five consecutive days of tamoxifen injection. (B) Insulin tolerance test and AUC of nine-month-old mice measured 19 days after five consecutive days of tamoxifen injection. (A and B) N=4 ($Bdh1^{+/+}$ male) and 5 ($Bdh1^{Thy1-/-}$ male) mice per group. Data are means ± SD. (C) Body weight, blood glucose, and plasma βHB levels of four- to five-month-old mice after seven days control diet (CD) or ketogenic diet (KD). Both diets were fed three days after five consecutive days of tamoxifen injection. (D) qPCR of genes involved in ketone metabolism in the brain. (C and D) N=4 ($Bdh1^{+/+}$ male CD), 4 ($Bdh1^{Thy1-/-}$ male CD), 4 ($Bdh1^{+/+}$ male KD), 6 ($Bdh1^{Thy1-/-}$ male KD), 4 ($Bdh1^{+/+}$ female CD), 3 ($Bdh1^{Thy1-/-}$ female CD), 6 ($Bdh1^{+/+}$ female KD), and 4 ($Bdh1^{Thy1-/-}$ female KD) mice per group. P = 0.0053 (Hmgcs2, male, KD, t = 3.78, df = 8), p = 0.0061 (Bdh1, male, CD, t = 4.13, df = 6), p = 0.0038 (Bdh1, male, KD, t = 4.02, df = 8), p = 0.0080 (Bdh1, female, CD, t = 4.27, df = 5), and p = 0.0087 (Bdh1, female, KD, t = 3.45, df = 8) by unpaired two-tailed t-test assuming Gaussian distribution. Data are means ± SD.

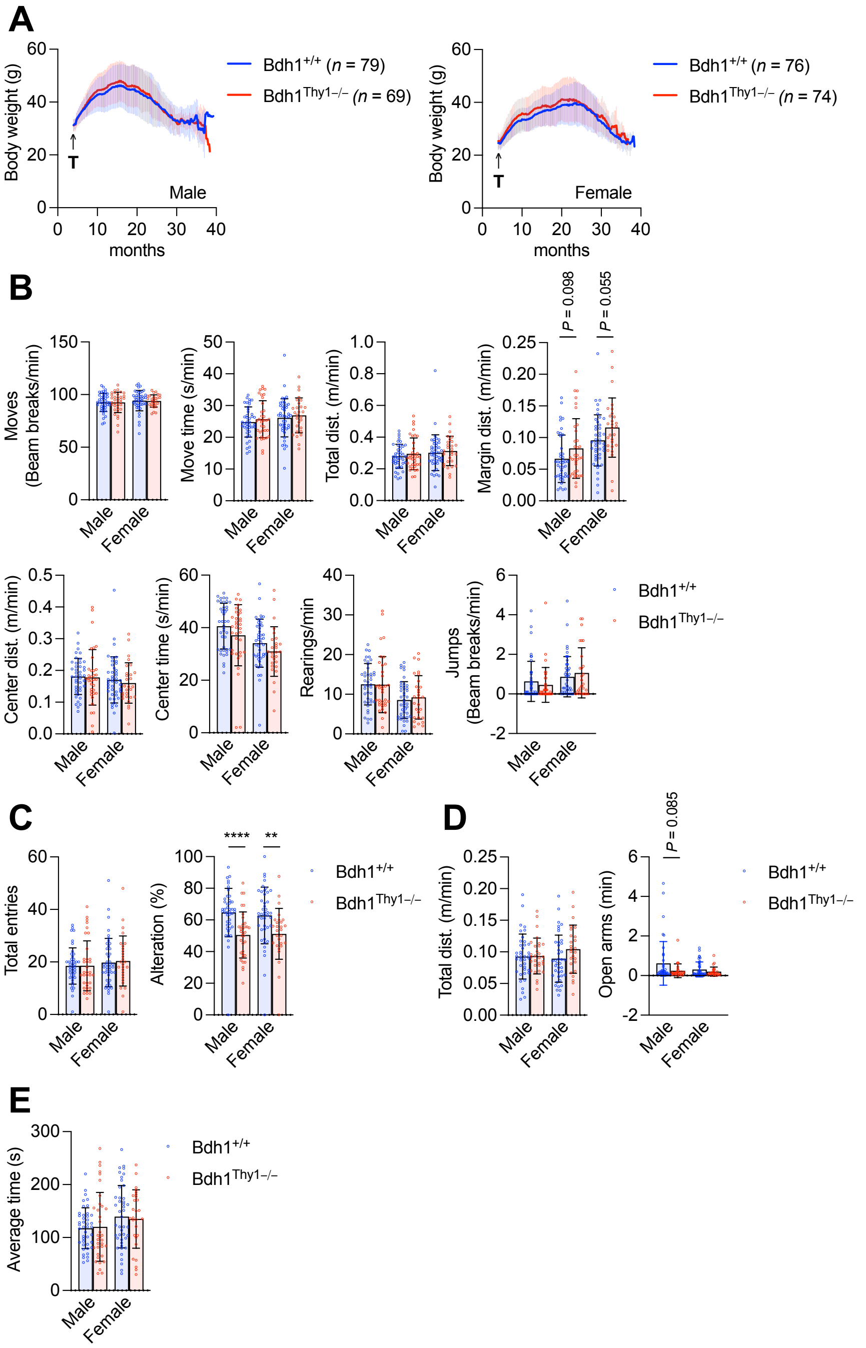


**Extended Data Fig. 5.**

(A) Body weight curves. (B) Open field test at 21 months old. N=42 (Bdh1$^{+/+}$ male), 35 (Bdh1$^{Thy1-/-}$ male), 44 (Bdh1$^{+/+}$ female), and 28 (Bdh1$^{Thy1-/-}$ female) mice per group. Data are means ± SD. (C) Y-maze at 21 months old. N=43 (Bdh1$^{+/+}$ male), 35 (Bdh1$^{Thy1-/-}$ male), 44 (Bdh1$^{+/+}$ female), and 28 (Bdh1$^{Thy1-/-}$ female) mice per group. $P < 0.0001$ (alteration, male, $t = 4.16$, $df = 76$) and $p = 0.0069$ (alteration, female, $t = 2.78$, $df = 70$) by unpaired two-tailed t-test assuming Gaussian distribution. Data are means ± SD. (D) Elevated plus maze at 24 months old. N=40 (Bdh1$^{+/+}$ male), 30 (Bdh1$^{Thy1-/-}$ male), 40 (Bdh1$^{+/+}$ female), and 26 (Bdh1$^{Thy1-/-}$ female) mice per group. Data are means ± SD. (E) Rotarod at 21 months old. N=43 (Bdh1$^{+/+}$ male), 35 (Bdh1$^{Thy1-/-}$ male), 44 (Bdh1$^{+/+}$ female), and 28 (Bdh1$^{Thy1-/-}$ female) mice per group. Data are means ± SD.

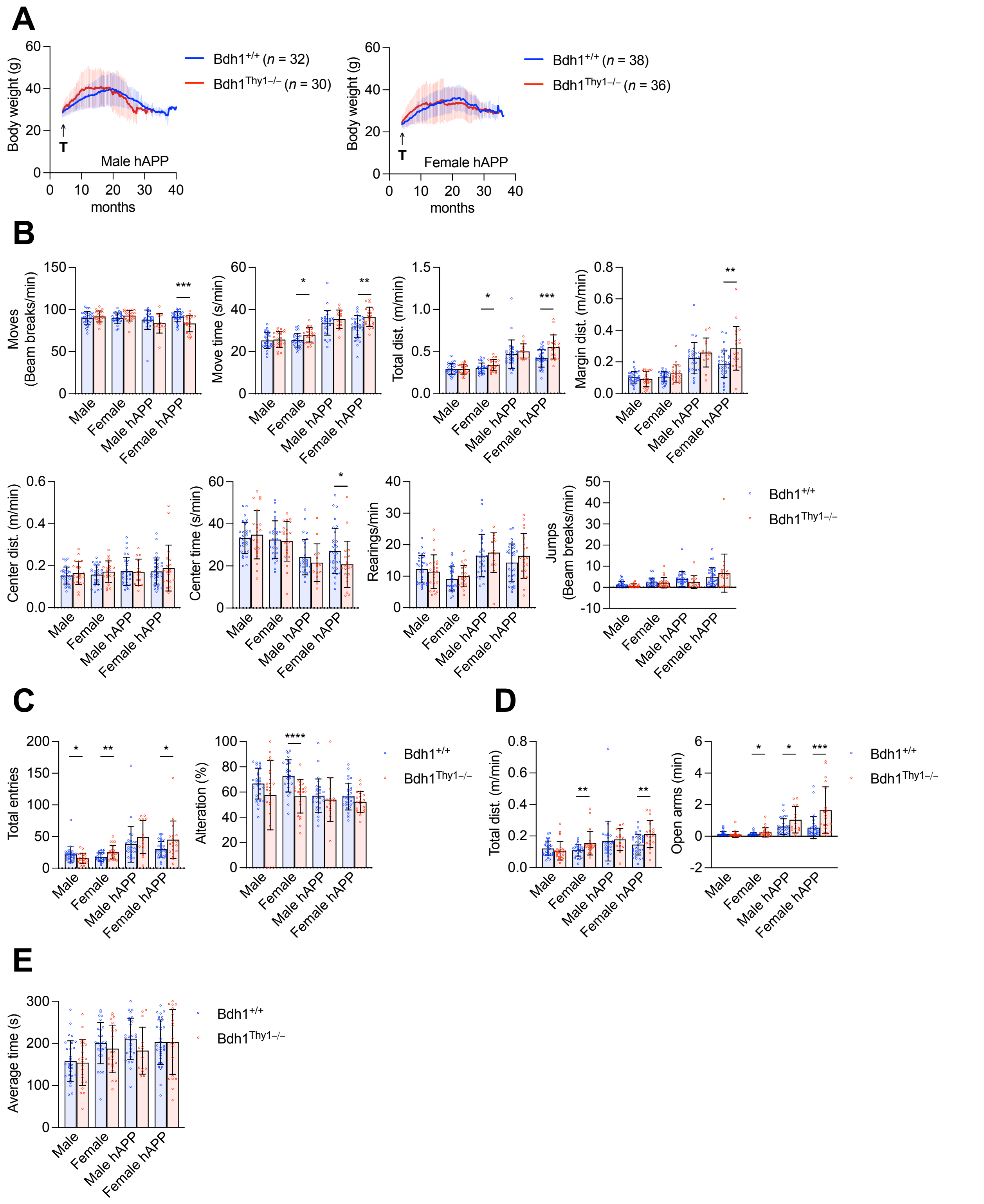


**Extended Data Fig. 6.**

(A) Body weight curves. (B) Open field test at 12 months old. N=30 (Bdh1$^{+/+}$ male), 23 (Bdh1$^{Thy1-/-}$ male), 27 (Bdh1$^{+/+}$ female), 22 (Bdh1$^{Thy1-/-}$ female), 28 (hAPPJ20/Bdh1$^{+/+}$ male), 14 (hAPPJ20/Bdh1$^{Thy1-/-}$ male), 33 (hAPPJ20/Bdh1$^{+/+}$ female), and 20 (hAPPJ20/Bdh1$^{Thy1-/-}$ female) mice per group. P = 0.00060 (moves, hAPPJ20 female, t = 3.66, df = 51), p = 0.011 (move time, female, t = 2.64, df = 47), p = 0.0020 (move time, hAPPJ20 female, t = 3.25, df = 51), p = 0.49

(total dist., female, t = 2.02, df = 47), p = 0.00024 (total dist., hAPPJ20 female, t = 3.95, df = 51), p = 0.0027 (margin dist., hAPPJ20 female, t = 3.16, df = 51), and p = 0.043 (center time, hAPPJ20 female, t = 2.07, df = 51) by unpaired two-tailed t-test assuming Gaussian distribution. Data are means ± SD. (C) Y-maze at 12 months old. N=30 (Bdh1$^{+/+}$ male), 23 (Bdh1$^{Thy1-/-}$ male), 27 (Bdh1$^{+/+}$ female), 22 (Bdh1$^{neu-/-}$ female), 28 (hAPPJ20/Bdh1$^{+/+}$ male), 14 (hAPPJ20/Bdh1$^{Thy1-/-}$ male), 33 (hAPPJ20/Bdh1$^{+/+}$ female), and 20 (hAPPJ20/Bdh1$^{Thy1-/-}$ female) mice per group. P = 0.046 (total entries, male, t = 2.05, df = 51), p = 0.0032 (total entries, female, t = 3.11, df = 46), p = 0.013 (total entries, hAPPJ20 female, t = 2.59, df = 50), and p < 0.0001 (alteration, female, t = 4.31, df = 46) by unpaired two-tailed t-test assuming Gaussian distribution. Data are means ± SD. (D) Elevated plus maze at 12 months old. N=30 (Bdh1$^{+/+}$ male), 23 (Bdh1$^{Thy1-/-}$ male), 27 (Bdh1$^{+/+}$ female), 22 (Bdh1$^{Thy1-/-}$ female), 28 (hAPPJ20/Bdh1$^{+/+}$ male), 14 (hAPPJ20/Bdh1$^{Thy1-/-}$ male), 33 (hAPPJ20/Bdh1$^{+/+}$ female), and 20 (hAPPJ20/Bdh1$^{Thy1-/-}$ female) mice per group. P = 0.0094 (total dist., female, t = 2.71, df = 47), p = 0.0022 (total dist., hAPPJ20 female, t = 3.23, df = 51), p = 0.043 (open arms, female, t = 2.08, df = 47), p = 0.042 (open arms, t = 2.01, df = 40), and p = 0.00060 (open arms, hAPPJ20, female, t = 3.66, df = 51) by unpaired two-tailed t-test assuming Gaussian distribution. Data are means ± SD. (E) Rotarod at 12 months old. N=30 (Bdh1$^{+/+}$ male), 23 (Bdh1$^{Thy1-/-}$ male), 27 (Bdh1$^{+/+}$ female), 22 (Bdh1$^{Thy1-/-}$ female), 28 (hAPPJ20/Bdh1$^{+/+}$ male), 14 (hAPPJ20/Bdh1$^{Thy1-/-}$ male), 33 (hAPPJ20/Bdh1$^{+/+}$ female), and 20 (hAPPJ20/Bdh1$^{Thy1-/-}$ female) mice per group. Data are means ± SD.
